# Adult stem cell characterization from the *Medial Gastrocnemius* and *Semitendinosus* muscles in early development of cerebral palsy pathology

**DOI:** 10.1101/2023.01.10.523388

**Authors:** M Corvelyn, J Meirlevede, J Deschrevel, E Huyghe, E De Wachter, G Gayan-Ramirez, M Sampaolesi, A Van Campenhout, K Desloovere, D Costamagna

## Abstract

Cerebral palsy (CP) is one of the most common lifelong conditions leading to childhood physical disability. Literature reported previously altered muscle properties such as lower number of satellite cells (SCs), with altered fusion capacity. However, these observations highly vary among studies, possibly due to heterogeneity in patient population, lack of appropriate control data, methodology and different assessed muscle.

In this study we aimed to strengthen previous observations and to understand the heterogeneity of CP muscle pathology. Myogenic differentiation of SCs from the *Medial Gastrocnemius* (MG) muscle of patients with CP (n=16, 3-9 years old) showed higher fusion capacity compared to age-matched typically developing children (TD, n=13). Furthermore, we uniquely assessed cells of two different lower limb muscles and showed a decreased myogenic potency in cells from the *Semitendinosus* (ST) compared to the MG. Longitudinal assessments, one year after the first botulinum toxin treatment, showed slightly reduced SC representations and lower fusion capacity. Finally, we proved the robustness of our data, by assessing in parallel the myogenic capacity of two samples from the same TD muscle.

In conclusion, these data confirmed previous findings of increased SC fusion capacity from MG muscle of young patients with CP compared to age-matched TD. Further elaboration is reported on potential factors contributing to heterogeneity, such as assessed muscle, CP progression and reliability of primary outcome parameters.

## Introduction

With an incidence of 1 in 500 live births, cerebral palsy (CP) is one of the most common lifelong conditions leading to childhood physical disability ^1^. It is considered to be originated from a neural lesion in the immature brain, leading to progressive musculoskeletal symptoms. Clinically, CP manifests itself on both neural and muscular level (spasticity, increased muscle stiffness and contractures, decreased strength and muscular control), resulting in decreased functional ability such as disturbed gait ^2–4^. These patients can be classified following the Gross Motor Function Classification System (GMFCS), from levels I to V, based on their functional abilities and limitations ^5,6^. Treatment mainly consists of management of the symptoms at the muscle level, including physiotherapy, orthoses, Botulinum Neurotoxin A (BoNT) injections and orthopaedic surgery ^4,7^. However, knowledge on the effects of treatment on CP muscle morphology is incomplete and totally lacking on cellular level ^4^. Hence, there is much interest in better understanding the underlying pathomorphology of skeletal muscles in patients with CP and thereby as well the effects of treatment ^3^.

Literature has reported altered muscle properties compared to typically developing (TD) muscles, mainly in adolescents and adults. Microscopically, increased collagen content and abnormal organization in the extra-cellular matrix have been reported and linked to muscle stiffness ^8,9^. Furthermore, some studies have reported smaller muscle fibers, varying predominance of muscle fiber type and increased adipogenic deposition ^10–13^. However, cells involved in these altered muscle properties and its onset and development are still poorly investigated.

Some groups have already shown a significant reduction in numbers of satellite cells (SCs) in contractured muscle of patients with CP ^14–16^. SCs are adult stem cells that get primarily involved in regeneration processes postnatally by fusion with damaged muscle fibers. These quiescent cells that are located between the sarcolemma and the basal lamina of the muscle, become activated to self-renew or to enter differentiation and express, amongst other transcription factors, MYOD and subsequently structural proteins such as MYOSIN HEAVY CHAIN (MyHC)^17,18^. Unfortunately, research on the functionality of these SCs in the muscles of patients with CP is scarce. Domenighetti et al. described lower fusion index (FI) values in CP-derived SCs (patients aged 3-18 years) and smaller myotubes compared to those of TD adolescents (aged 14-18 years), based on hamstring biopsies ^19^. Our group previously reported higher FI values and larger myotubes based on MyHC+ areas of SCs from younger patients with CP (aged 3-9 years) derived from microbiopsies of the *Medial Gastrocnemius* (MG) compared to aged-matched TD children ^20^. Potential reasons for these conflicting results, such as differences in isolation material and protocol, patient diversity and the variety of the assessed muscle, have already been suggested and need to be further explored to improve the understanding of the heterogeneity of the CP muscle pathology, before drawing generalized conclusions on muscle stem cell alterations in CP ^20^. In particular, Lieber and Domenighetti raised concerns on the purity and *in vitro* fusion capacity of myoblasts from young CP and age-matched TD, published by our group ^21^. Unfortunately, a proper reference for *in vitro* SC differentiation at these young ages is still missing or poorly reported. Therefore, to shed a light on these controversies and to broaden our previous findings, we corroborated our TD sample size and better characterized the purity and myogenic potential from microbiopsy-derived SCs ^20^. Furthermore, as the muscle-specificity could potentially partly explain the observed heterogeneity in the fusion potential of SCs, a logic next step was to isolate and characterize stem cells from different lower limb muscles. Lastly, more knowledge on the reliability of stem cell characterization parameters based on repeated assessments could further strengthen the value of the study results.

In addition to SCs, also mesoangioblasts (MABs) have been recognized for their role in regeneration. Next to SCs, MABs are also able to directly fuse with damaged muscle fibers. In case of muscular dystrophies, MABs have shown to reconstitute the SC pool in case SCs were exhausted or insufficient for muscle regeneration ^22–24^. Furthermore, also fibro-adipogenic progenitor cells (FAPs) have shown to be involved in the muscle regeneration processes, firstly by synthesizing the transient connective tissue necessary to maintain the structural integrity during regeneration and secondly by supporting myogenesis/the myogenic process by stimulating SC activation ^25–27^. However, in cases of chronic injury, these cells start to highly proliferate and could differentiate towards adipocytes leading to adipogenic muscle loss ^26,28^. Unfortunately, data on the role of these progenitor cells in the light of CP muscle pathology are almost completely lacking and did not show altered differentiation capacity in young children with CP ^20^.

Additionally, for previous described microscopic CP alterations, biopsies were usually obtained during invasive surgical procedures, such as single event multilevel orthopaedic surgery. As these procedures are commonly scheduled for children older than 8-10 years, data represent findings for a limited subgroup of the patient population, especially lacking younger subjects. Moreover, including ideal age-matched typically developing subjects, without conditions affecting the assessed muscle, has been proven to be challenging ^16,19,29^. Furthermore, the heterogeneity of available data is further increased by the different muscles (in case of the lower limb muscles; *Gastrocnemius, Soleus, Semitendinosus, Gracilis*, etc) considered in previous studies ^3,30^. The *Medial Gastrocnemius* (MG) muscle is quite commonly assessed, as this muscle is frequently involved in children with CP, leading to pathological ankle motions in gait and is rather large and accessible ^31^. The group of hamstring muscles, which has a more prominent role in posture, has been investigated in multiple studies as well, without discriminating between the different muscles of the hamstrings, i.e. the *Semitendinosus* (ST) ^11,19,32^. Finally, to assess adult muscle stem cell properties distinct methods (i.e. open biopsies and cell analysis after enzymatic digestion or muscle (micro-)biopsies followed by cell amplification *in vitro*) have been applied, leading to varying levels of *in vitro* potencies ^19,20^. Therefore, care should be taken to generalize conclusions, on which potential new therapeutic agents could be tested. To improve knowledge on the muscle pathology and its evolution, more research is thus needed on biopsies from different muscles, and especially collected in young children with CP, including appropriate age-matched TD children. These novel data can improve the understanding of the complex nature of intrinsic muscle impairment in CP, which may eventually help to optimize treatment.

Finally, while first attempts in characterizing stem cell features of patients have been made, information on the progression of these features and on their role in treatment is still not available. Since previous studies mainly applied invasive muscle biopsy methods, no longitudinal assessments have been possible. Yet, these longitudinal data are clinically relevant. For example, muscle imaging and animal studies highlighted that BoNT injections that are planned to reduce spasticity may induce atrophy and increase the fibro/fatty content in the muscle ^4,33,34^. Though, the effects of BoNT on microscopical level or on the stem cells of the muscle are much less studied, but are required for optimizing treatment.

Driven by the increasing indications of stem cell involvement in the altered muscle features of patients with CP ^19,20,35^, the current study addressed a series of uncertainties that may help to understand the heterogeneity in CP muscle pathology, especially with respect to muscle stem cell behaviour. Thereby we defined three objectives for this study: confirming previous reported CP SC features by increasing TD sample size, assessing SC differences between muscles or over time and investigating multiple potential involved stem cell populations. The first objective was to analyze an increased dataset of muscle microbiopsies from age-matched TD subjects to confirm and generalize our earlier preliminary findings on SC behaviour ^20^. Thereby, a more thorough characterization of the obtained stem cells was presented to specifically address the above mentioned two major concerns recently indicated by Lieber and colleagues ^21^. The second objective was to investigate potential factors leading to heterogeneity of muscle stem cell features. Thereby, SCs were collected and assessed from both the MG as well as the ST muscles of the same patients, to better understand the muscle-specificity of the observed phenotype. Additionally, using the muscle microbiopsy technique, a unique longitudinal preliminary assessment of CP-derived stem cell properties of the same muscle was performed. This allowed first insights into the progression of CP pathology on the level of the muscle stem cells and into the exploration of possible effects caused by previous BoNT treatment. Furthermore, for a better understanding of these longitudinal data and understanding the robustness of the primary outcome parameters, repeatability experiments were performed to quantify and decrease the risk of misinterpretation of potential effects due to the procedure or to the biopsy quality. The third objective was to examine the role of MABs, FAPs and interstitial cells (ICs) in the observed muscle alterations in both the MG and ST, cross-sectionally as well as longitudinally, i.e. prior to- and post-BoNT treatment.

## Materials and methods

### Muscle microbiopsy collection

This study protocol was approved by the Ethical Committee of the University Hospitals of Leuven, Belgium (S61110 and S62645). Written informed consent was provided by the parents. Children with cerebral palsy (CP) were recruited from the CP Reference Centre, whereas typically developing (TD) children were recruited from the Traumatology Unit for upper limb surgeries, or from the Ear, Nose and Throat Unit for other procedures at the University Hospitals Leuven (Belgium). A group of 17 patients with CP (age 3-9 years; Gross Motor Function Classification System levels I-III) and 13 age-matched TD children were included in this study (see Table 1). In the CP group, children with presence of dystonia or ataxia, BoNT injections within the last 6 months, orthopedic surgery less than 2 years before data collection, as well as any previous muscle surgery on the MG or ST were excluded. Additionally, TD children were further excluded when they had a history of neurological problems or when they were involved in a high-performance sporting program. All biopsies were collected during interventions requiring general anaesthesia related to orthopaedic interventions (including BoNT-injections). Combined microbiopsies from the muscle mid-belly of the MG and of the ST in the same child were obtained for a subgroup of 6 CP and 3 TD children. Four of the enrolled children with CP were included for longitudinal assessment, for that purpose a first biopsy of the MG was taken when they were BoNT-naïve at the time of study enrolment, and a second one was collected over one year later (13 to 19 months after first BoNT treatment). Unfortunately, sample size for these longitudinal studies was small as the recruitment was hampered due to the covid-situation. Two muscle biopsies for parallel stem cell assessments were obtained at the same time from the MG muscle of two TD children for the repeatability investigation of the primary outcome parameters. The biopsy collections were performed percutaneously under ultrasound guidance, with a microbiopsy needle (16-gauge, Bard). Clinical tolerance was good (based on questionnaire for parents two days post-biopsy). Furthermore, a questionnaire about the hobbies and intensity levels of activity was used for all enrolled subjects. Even though the primary goal of this study was to investigate muscle stem cell features for which the collected biopsies were entirely used, for a limited number of the enrolled patients, we could collect additional microbiopsies during the same session. These allowed explorative histological assessment of the muscle tissue via Haematoxylin and Eosin staining, as well as prospective *ex vivo* SC counting, to further describe the muscle and complement the *in vitro* cell culture data for a small subset of the participants.

**Table 1.**
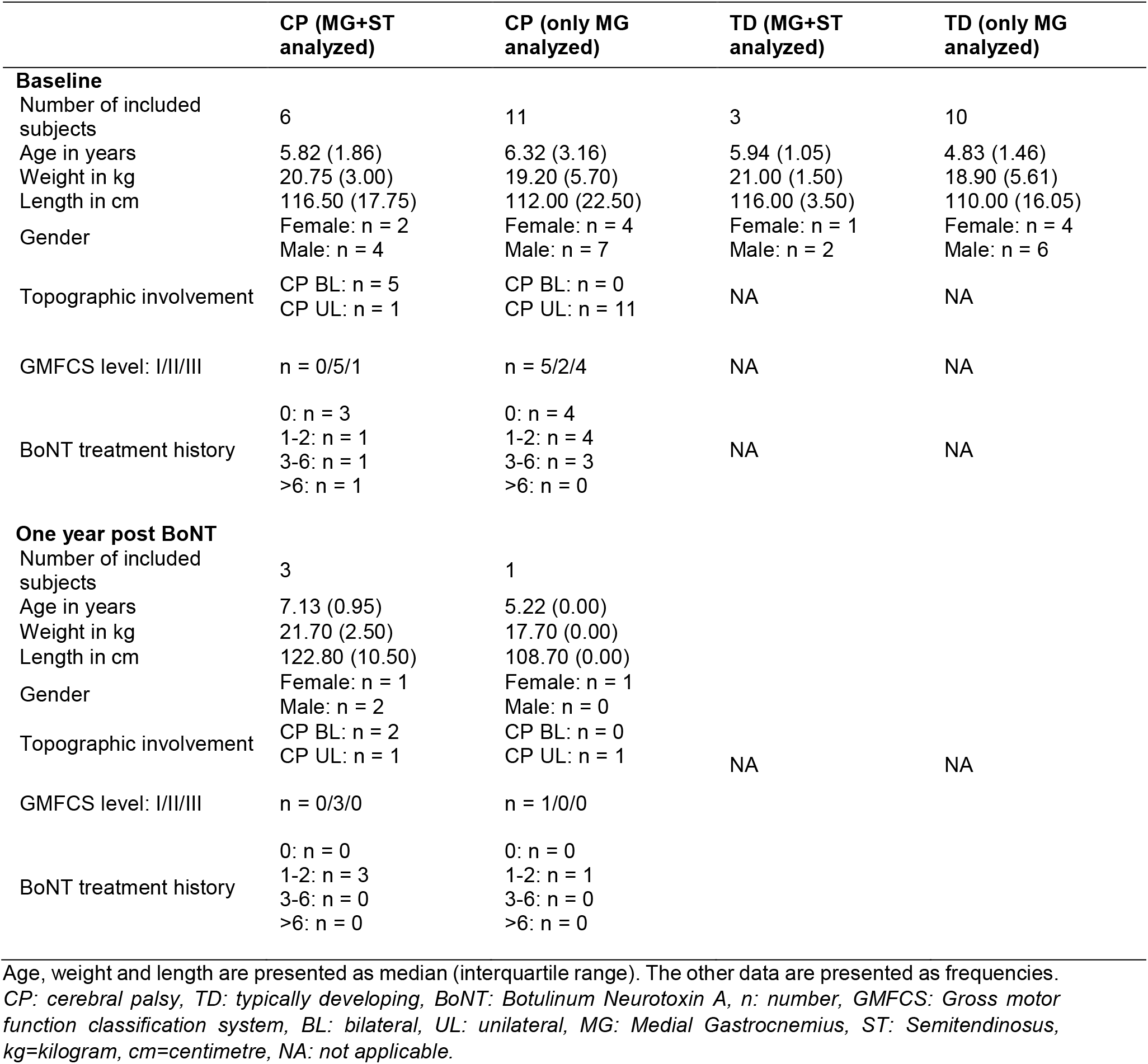
Demographic and anthropometric data for recruited subjects.

### Study design

For each of the three study objectives, specific analyses were planned on specific muscle samples of the entire study group or on a subgroup of the enrolled subjects. Table 2 gives an overview of these analyses, the available datasets, along with their main outcome parameters, structured per objective. More details on each analysis are described in the subsequent sections.

**Table 2.** Study design based on three objectives.

|  | Included patients | Muscle | Techniques | (Primary) outcomes |
| --- | --- | --- | --- | --- |
| **Objectives** |  |  |  |  |
| **1) Satellite cell alterations** |  |  |  |  |
| TD vs CP | TD: n = 13<br>CP: n = 16 | MG | IF stainings, Flow cytometry | **SC Fusion index (Day 6)** |
| **2) Satellite cell heterogeneity** |  |  |  |  |
| 2.1 MG vs ST | TD: n = 3<br>CP: n = 6 | MG and ST | FACS, IF stainings<br>Muscle section analysis | FACS fractions, KI67 and MYOD expression, **Fusion index (Day 6)** |
| 2.2 Longitudinal (Pre-post BoNT) | CP: n = 4 | MG | FACS, IF stainings | FACS fractions, MYOD expression<br>**Fusion index (Day 6)** |
| 2.3 Repeatability | TD: n = 2 | MG | FACS, IF stainings | FACS fractions, MYOD expression<br>**Fusion index (Day 6)** |
| **3) Supportive stem cell characteristics (MAB, FAP, IC)** |  |  |  |  |
| 3.1 MG vs ST | TD: n = 3<br>CP: n = 5 | MG and ST | FACS, IF stainings<br>Muscle section analysis | FACS fractions, MYOD expression, **Fusion index (Day 6)** |
| 3.2 Longitudinal (Pre-post BoNT) | CP: n = 4 | MG | FACS, IF stainings | Oil Red O quantification<br>FACS fractions, **Fusion index (Day 6)** |

*TD: typically developing, CP: cerebral palsy, BoNT: Botulinum Neurotoxin Type A, n: number, GMFCS: Gross motor function classification scale, MG: medial gastrocnemius, ST: semitendinosus, SC: satellite cell, MAB: mesoangioblast, FAP: fibro-adipogenic progenitors, IC: interstitial cell, IF: immunofluorescence, FACS: Fluorescence activated cell sorting. Primary outcome parameters are indicated in bold.*
Footnote Table 2: As specified above, the number of participants per subgroup, structured per objective varied. These subgroup sample sizes were primarily defined by practical reasons, such as ethical restrictions on the number of biopsies per patient and the time and costs of analyses. Furthermore, the impact of the covid-19 pandemic led to an unfortunate loss of patients being recruited for longitudinal assessments after being enrolled for a first biopsy collection at a BoNT-naïve state. Hence, we believe that the selected subgroups were unbiased.

### Cell culture and stem cell isolation via FACS

Cells were amplified and isolated according to a previous published protocol ^20^. Cell populations were isolated by serial fluorescent activated cell sorting (FACS) using a BD FACSAria II (BD biosciences) or Sony MA9000 (Sony) device. Calcein violet (10µM/1 × 10^6^ cells, eBioscience), was used as additional viability control. Firstly, satellite cell-like progenitors (SCs as for the previous data set (16)), were sorted based on CD56 marker ^36^. CD56 negative cells were further amplified and sorted in order to isolate mesoangioblasts (MABs), based on the presence of ALKALINE PHOSPHATASE (CD56-ALP+ PDGFRa-cells), while fibro-adipogenic progenitor cells (FAPs) were sorted for PLATELET DERIVED GROWTH FACTOR RECEPTOR ALPHA (CD56-ALP-PDGFRa+ cells). Interstitial cells (ICs) were collected based on the absence of these markers (CD56-ALP-PDGFRa-). All antibody details are specified in supplementary table 1. Based on the available literature on these cell types and to facilitate the clarity of the paper, we will simplify referring to these progenitor populations as SCs, MABs, FAPs and ICs. All cell populations underwent further proliferation and differentiation assays followed by immunofluorescent staining as further described.

### Stem cell characterization by flow cytometry analysis

Antibody titrations for FACS optimization and further flow cytometry analyses were performed with a FACSCanto II HTS (BD biosciences). Experiments were performed to prove that the obtained cell population before FACS isolation was pure, based on stemness marker CD34, haematopoietic marker CD45 (and endothelial markers CD144 and CD31. Positive control samples, U266 multiple myeloma cell line and iPSC-derived endothelial cells, were obtained to show the specificity of the antibodies. Data were analyzed with FlowJo v10.6.1 software.

### *In vitro* differentiation assays

Myogenic (SCs and MABs) and adipogenic (MABs, FAPs and ICs) differentiation assays were performed as previously reported ^20^. Cells included for proliferation analyses were seeded at a density of 3 × 10³ cells/cm². In the light of the repeatability assessments, a subgroup of SC cells (n = 5) underwent myogenic differentiation at different passages (P and P+2).

### Immunofluorescent staining and imaging of differentiated cells

Cells were cultured in 96 well dishes (Thermo Fisher Scientific) and fixed with 4% paraformaldehyde. Immunofluorescent (IF) staining was performed according to our previously published protocol ^20^. All used primary antibodies are listed in supplementary table 1. Appropriate secondary antibodies were used (1:500, Alexa Fluor® donkey 488 or 594, Thermo Fisher Scientific). Hoechst (1:3000 in PBS, Thermo Fisher Scientific) was used for indicating nuclei. Fusion index (FI) was calculated as ratio between at least 2 myonuclei per MYOSIN HEAVY CHAIN (MyHC) positive area and the total number of nuclei in a field of view. Three randomized images per well have been analyzed per assessment. Visualization occurred with an Eclipse Ti Microscope (Nikon) and NIS-Elements AR 4.11 software or an inverted DMi8 microscope (Leica) and LASX software, depending on the staining.

### Oil Red O staining and quantification of adipogenic differentiation potential

After 10 days of adipogenic differentiation, Oil Red O (ORO) staining was performed to assess the deposition of lipid droplets, as previously described ^20^. Cells were incubated for 50 minutes with ORO solution (65% of 0.5% w/v Oil Red O in isopropanol (Thermo Fisher Scientific) and MQ water). Secondly, the previously described standard IF protocol for cells was applied for visualization of the adipocyte membrane with PERILIPIN. ORO was measured by extracting lipids with a petrol ether/isopropanol mixture (3:2) and quantified for their absorbance at 490 nm for 0.1 s with Victor Spectrophotometer (PerkinElmer). Standard curve was applied and quantification was expressed in µg of pure ORO powder.

### Histological and immunofluorescent analyses on muscle sections

As a side trajectory, for a subgroup of patients, we included additional microbiopsies to perform histological assessment of the muscle tissue, which allowed to complement the *in vitro* cell culture data and assess the CP muscle in an explorative manner. Muscle microbiopsies were snap-frozen in isopentane cooled in liquid nitrogen and stored at −80°C. Five μm muscle sections were obtained using a CryoStarTM NX70 Cryostat (Thermo Fisher Scientific) while kept at −20°C. Haematoxylin and Eosin staining was performed on few ST muscle sections by incubating the slides at RT for 1 min in hematoxylin solution (Merck). Slides were then washed for 10 min in running tap water before incubating for 1 min in eosin solution (Sigma-Aldrich). Slides were then washed extensively in tap water. To dehydrate, sections were incubated consecutively for 10 sec in 70%, 90% and 100% ethanol and 5 min in xylene/ethanol (1:1, Merck) and 5 min in xylene. Muscle sections were dried, before mounting with DPX mountaint (Merck) and qualitatively assessed for contractile material and extra-cellular matrix.

IF staining for SC counting on MG and ST muscle slides was performed based on co-localization of PAX7 and Dapi. Sections were placed at room temperature (RT) for 10 min in a humid chamber followed by 10 min fixation in cold acetone. After washing with PBS, slides were incubated with 10% goat serum (Invitrogen) for 1 h. Primary antibodies were applied in blocking solution overnight at 4°C. After 3 × 5 min washing with PBS, muscle slides were incubated for 1 h at RT with appropriate secondary antibodies (1:500, Alexa Fluor® goat 488 or 680, Thermo Fisher Scientific). After 3 × 5 min washing with PBS, Dapi (1:50 in PBS, Thermo Fisher Scientific) was added for 1 min and the samples were extensively washed with PBS and mounted with ProLong® Gold antifade reagent (Molecular Probes). Visualization was performed with an inverted DMi8 microscope (Leica) and LASX software.

### Data analyses

Data in this study are represented using means ± standard deviation. Boxplots have been used in the visualization of FI values, percentages of KI67+ or MYOD+ nuclei and ORO quantifications. The boxplots indicate the 25th to 75th percentiles, while the whiskers indicate the minimum and maximum values. The centre of the boxplot indicates the median. FACS fractions are indicated by individual values and the bar represents the mean. Unpaired T-tests were performed for comparison between the TD and CP groups (two-tailed, p < 0.05). Paired T-tests were applied to compare data from the MG and ST muscles as well as for longitudinal comparisons. Data were pooled from CP and TD children, when relevant, and only in case no differences were found between groups. Repeatability assessments were explored in a qualitative way. Analyses of SC FI values with a difference of two passages was performed using a Two-tailed paired T-test. Numbers of analyzed data are always indicated in the results section as well as in the figure legends. Statistical analysis and simple linear regression analysis were performed using GraphPad Prism 8 software. Significance of the differences was reported using ‘$’ for TD and CP, ‘*’ for comparison between MG and ST and ‘§’ for assessment prior to and after BoNT treatment and for all of them it represents p-values smaller than 0.05.

## Results

### Satellite cell alterations of patients with CP compared to TD (objective 1)

Based on immunofluorescence (IF) images for MYOSIN HEAVY CHAIN (MyHC) and a nuclear dye, Hoechst, we found higher fusion index (FI) values for satellite cells (SCs) derived from *Medial Gastrocnemius* (MG) muscle of children with CP (41.60 ± 17.94%; n = 16) compared to those of TD (28.33 ± 14.39%; n = 13; $ p < 0.05; Figure 1). These data were added to previous published results and implemented with additional TD data to increase power ^20^.

**Figure 1.**
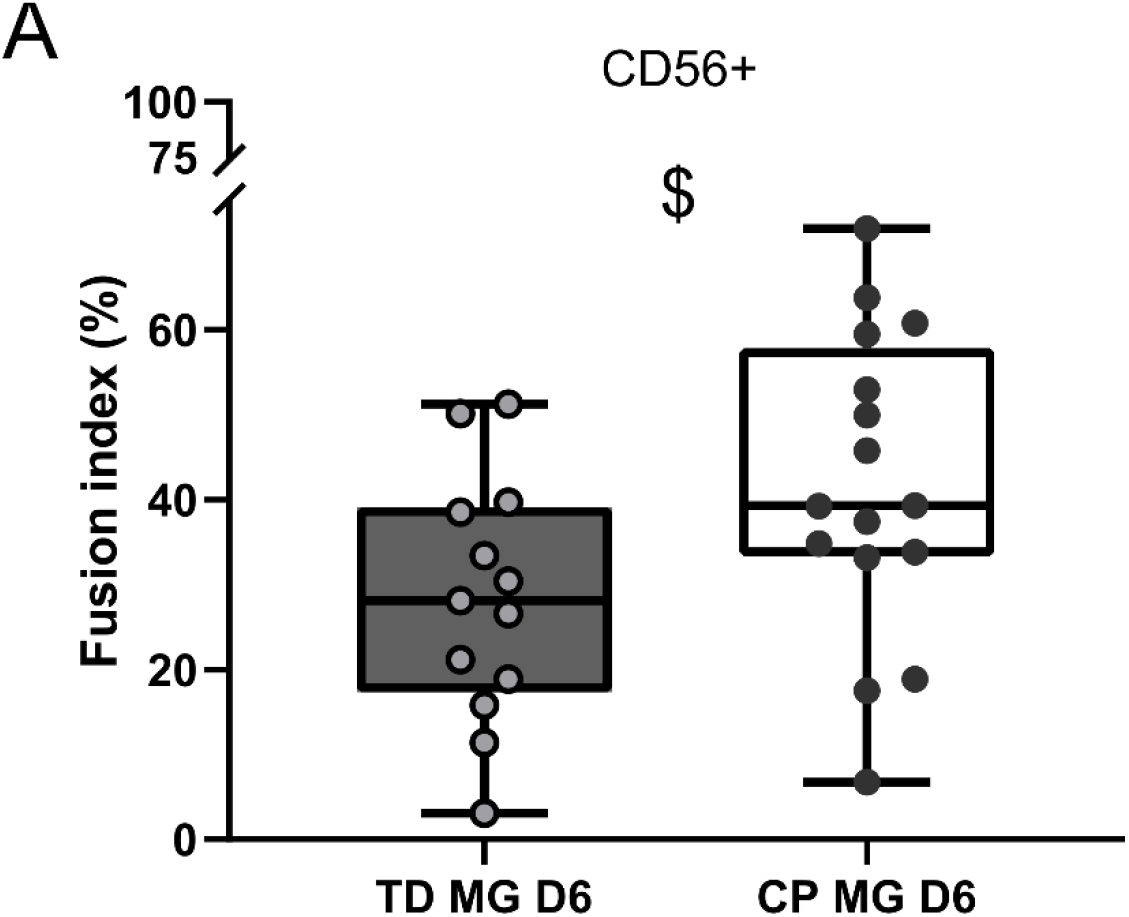
Fusion index of satellite cells (CD56+) from the *Medial Gastrocnemius* (MG) muscle. **(A)** Fusion indexes of SC myogenic differentiation at day 6, completed with additional data from both typically developing children (TD) and patients with cerebral palsy (CP). Data shown by boxplots, every dot represents an individual subject. TD: n = 13, CP: n = 16, $ p < 0.05

Flow cytometry was performed on the total population of cells extracted from the muscle to quantify additional markers such as markers of stemness (CD34) or markers that are highly expressed by blood cells (CD45) or endothelial cells (CD31 and CD144, TD and CP: n = 6). Analysis showed that on average 1.78 ± 1.97% of all cells were positive for CD34 marker in TD and 0.69 ± 0.88% in CP. On average 0.28 ± 0.30% of cells derived from TD samples and 0.05 ± 0.06% from CP were positive for CD45 haematopoietic marker. The endothelial markers CD144 and CD31, were expressed by 0.03 ± 0.07 % and 0.03 ± 0.06% of TD cells and 0.01 ± 0.01% and 0.03 ± 0.03% of CP-derived cells, respectively (Supplementary figure 1A, B). Positive control sample for haematopoietic cells, U266 lymphocytes, showed 11.5% CD45+ cells and 44.9% CD31+ cells (Supplementary figure 1C). Positive control sample for endothelial cells, hiPSC-derived endothelial cells were included and showed expression of markers CD144 by 8.7% and CD31 by 84.8% (Supplementary figure 1D). Both control samples showed expression levels according to previous literature (34,35). Overall, we showed higher fusion capacity of the SCs derived from the MG of patients with CP compared to those of TD without the interference of haematopoietic and endothelial cells.

### Satellite cell heterogeneity based on muscle-specificity (objective 2.1)

Firstly, in order to examine the observed heterogeneity amongst different studies with regard to muscle stem cell behaviour, we have assessed SCs from two different lower limb muscles (the MG and the *Semitendinosus* (ST)) of the same subjects. FACS analyses showed proportions of CD56+ cells, consequently referred to as satellite cells (SCs), of 38.8 ± 10.78% in TD (n = 3) and 34.05 ± 19.53% in CP (n = 6) for the MG muscle. In the ST muscle, SC were represented for 31.33 ± 13.26% in TD children and 25.38 ± 15.92% in CP (Figure 2A). Since no differences could be found between TD and CP, we pooled the data of both groups as we observed a slight reduced SC fractions in the ST muscle compared to MG (−8.27%), but this observation failed to reach statistical significance. Additionally, we quantified *ex vivo* SC representations based on PAX7+ nuclei in muscle sections, as this staining marks quiescent SCs. Correlation assessment of FACS fractions and *ex vivo* SC localization in muscle sections showed no correlation (R²=0.006, n = 12 (pooled TD, CP, MG and ST data); Supplementary figure 2).

**Figure 2.**
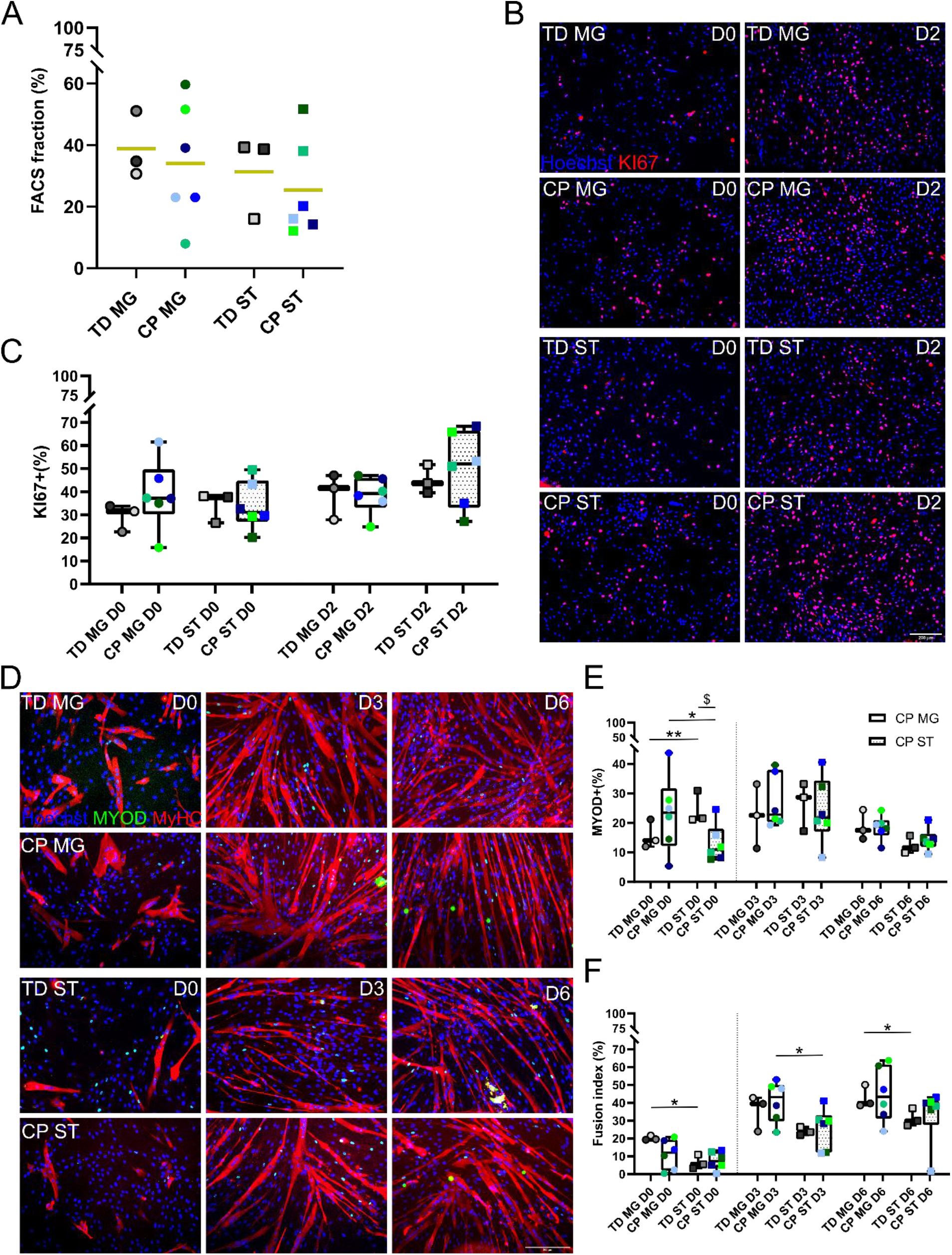
Satellite cell features of the *Medial Gastrocnemius* (MG) and *Semitendinosus* (ST) muscle in TD and CP children. **(A)** Fraction of obtained stem cells through FACS with CD56 marker, referred to as satellite cells (SCs). Data expressed as percentage of total sorted cells. The bar represents the mean. Dots (MG) and squares (ST) represent individual subjects indicated in different colours. **(B)** Representative immunofluorescent (IF) images from SCs derived from TD children and patients with CP for both MG and ST muscle at day 0 and day 2 during proliferation. KI67 is shown in red. Nuclei (blue) are counterstained using Hoechst. **(C)** Percentage of KI67 positive cells based on IF images. Data shown by boxplots, dots (MG) and squares (ST) represent individual subjects indicated in different colours. **(D)** Representative IF images from SCs derived from TD children and patients with CP for both MG and ST muscle at days 0, 3 and 6 of myogenic differentiation. MYOD (green) and MYOSIN HEAVY CHAIN (MyHC; red) are shown. Nuclei (blue) are counterstained using Hoechst. **(E)**. Percentage of MYOD+ cells based on IF images represented by boxplots for SCs from both TD children and CP patients. Dots (MG) and squares (ST) represent individual subjects indicated in different colours. **(F)** SC fusion indexes shown by boxplots. Dots (MG) and squares (ST) represent individual subjects indicated in different colours. Scale bars are 200 µm. TD: n = 3, CP: n = 6, * or $ p < 0.05, ** p < 0.01

SCs (CD56+) were analyzed for the expression of KI67 based on IF staining at day 0 and day 2 during proliferation conditions (Figure 2B, C). No differences between TD and CP nor between both muscles, were found on both time points. SCs underwent myogenic differentiation and were analyzed for MYOD expression and fusion index (FI) at days 0, 3 and 6 of differentiation (Figure 2D-F). Firstly, at day 0, the percentage of MYOD+ nuclei for the ST was significantly higher in the TD compared to the CP group (*p < 0.05). For the TD group, higher percentages of MYOD+ nuclei were found for the ST compared to the MG (24.56 ± 5.57% and 15.72 ± 4.84%; n = 3; **p < 0.01) at day 0, while the opposite was found for the CP group (ST: 13.01 ± 6.40% and MG: 23.06 ± 12.98%; n = 6; *p < 0.05). No differences were found between MYOD expression of both groups and muscles at days 3 and 6. Secondly, FI analysis showed no significant differences in the ability of these SCs to fuse into myotubes between TD and CP at all time points. Furthermore, FI values significantly increased in MG compared to ST at days 0 (MG: 20.17 ± 1.31%; ST: 6.57 ± 3.95%) and day 6 (MG: 42.78± 6.36%; ST: 31.42± 4.96%) for TD (n = 3; *p < 0.05). This increased FI was also reported at day 3 for CP (MG: 40.61 ± 11.43% and ST: 25.62 ± 11.49%; n = 6; *p < 0.05). As no significant differences were primarily present between TD and CP in this dataset, we secondarily pooled data of the MG and ST from all subjects (TD and CP, n = 9) to better explore potential muscle-specificity. SCs derived from the MG muscle had higher FI values compared to those of ST both at days 3 (MG: 38.87 ± 10.67% and ST: 25.08 ± 9.19%; p = 0.010) and day 6 (MG: 44.14 ± 12.75% and ST: 32.66 ± 12.65%; p = 0.012), and showed a trend towards higher FI at day 0 (p = 0.064). Taken together, we found higher fusion capacity of SCs derived from the MG compared to the ST for both the TD as well as the CP group during myogenic differentiation.

### Longitudinal effects on stem cell features from the MG of patients with CP (objective 2.2)

Secondly, to examine possible effects of the Botulinum Neurotoxin A (BoNT) treatments and/or the progression of the CP pathology on the muscle stem cells, four patients with CP were enrolled twice for assessing MG microbiopsies, prior to BoNT treatment (BoNT-naïve) and post BoNT treatment (BoNT treated). No significant differences in SC FACS fractions in patients with CP were observed comparing time points prior to and after BoNT treatment, although an average decrease of 16.4% was found (Figure 3A). Myogenic capacity of these SCs indicated that three out of 4 cases showed lower FI values after BoNT treatment, while in one case, FI was slightly higher at day 6 (Figure 3B-D). In general, SC FI values decreased on average with 9,71% in comparison to the pre-BoNT time point after 6 days of myogenic differentiation.

**Figure 3.**
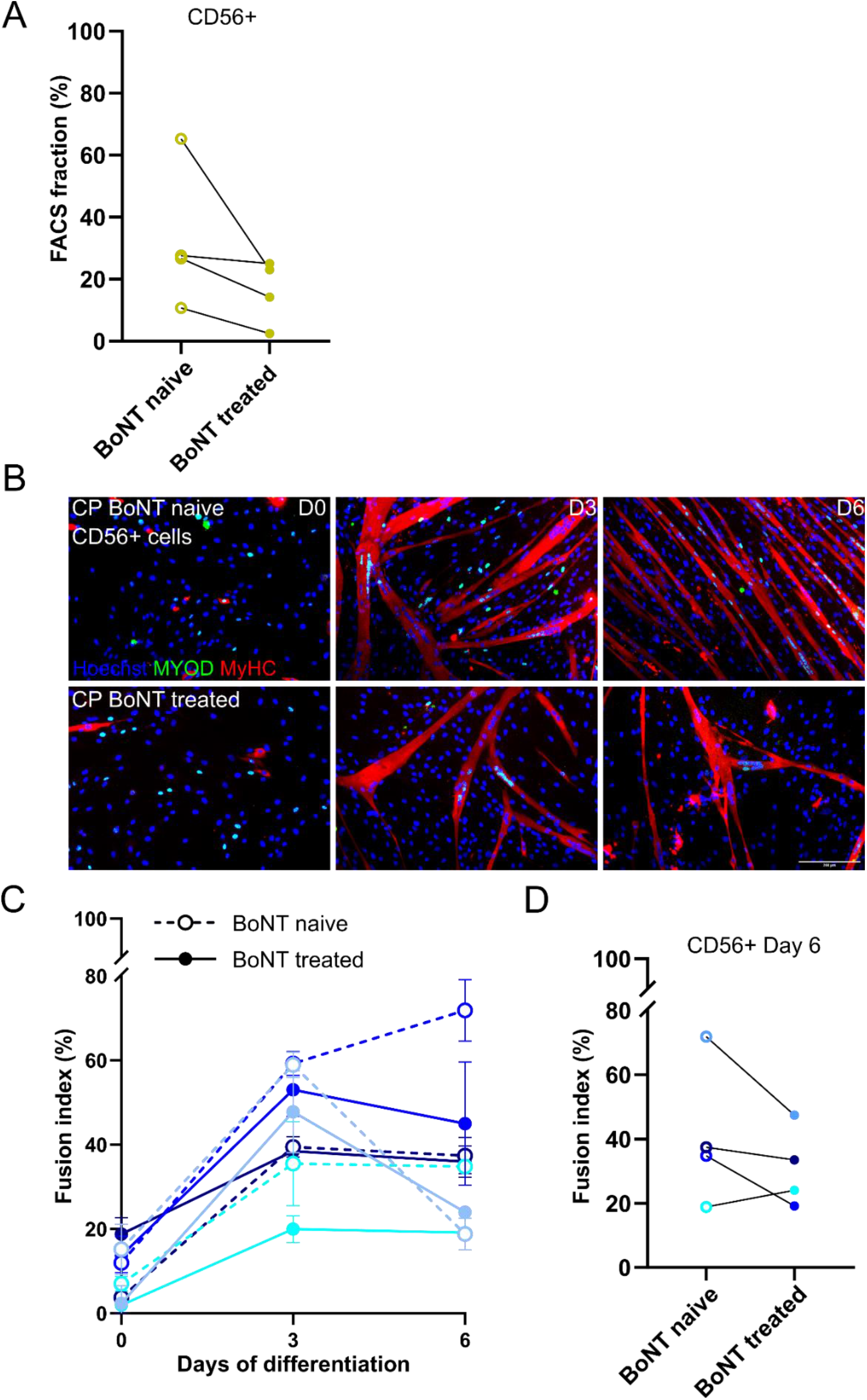
Satellite cell features prior to (BoNT naïve) and post Botulinum toxin treatment (BoNT treated). **(A)** Quantification of SCs (CD56+) from the *MG*, expressed in percentage of total sorted cells from patients with CP prior to (BoNT naïve) and post (BoNT treated) BoNT treatment. (n = 4) **(B)** Representative IF images of CP patient-derived SC myogenic differentiation at days 0, 3 and 6 from a BoNT naïve patient and at one year after BoNT treatment. MYOD (green) and MyHC (red) and nuclei (blue) are shown. Scale bar = 200 µm. **(C)** Fusion indexes of SCs from patients with CP at days 0, 3 and 6 of myogenic differentiation. Each subject is indicated in a different colour. FI of SCs from BoNT naive patients with CP are indicated by dotted lines and by solid lines after BoNT treatment. **(D)** Longitudinal effect on FI of SCs at day 6. Data from the same subject are connected through a black line from BoNT naive to BoNT treated.

Repeatability and reliability of the assessed outcome parameters (objective 2.3) Thirdly, to explore to what extent the observed findings could be influenced by the procedure or biopsy quality, two separate microbiopsies of the MG from TD children were obtained at the same time and at the same muscle location and processed in parallel (n = 2). FACS fractions for the CD56+ population fluctuated by 24.23% between repeated assessments (Supplementary figure 3A). For the myogenic parameters, percentages of MYOD+ nuclei and FI values remained stable at days 3 and 6, but showed rather high heterogeneity at day 0 (Supplementary figure 3B, C). On average a minor difference between repeated assessments of 4.66% was observed at day 3 and 1.59% at day 6 for MYOD expression, while FI differed with 2.45% and 7.46% at days 3 and 6, respectively.

In this light, we determined the effect of these SC FACS fluctuations on the FI measured at day 6 of myogenic differentiation by simple linear regression on the full dataset, including both TD (n = 13) and CP (n = 16) data (Supplementary figure 3D). There was no correlation between SC FACS fractions and the FI measured at day 6 of myogenic differentiation (R² = 0.1781). Furthermore, the effect of variance in passage number of SCs at the time of myogenic differentiation was assessed as this could vary with maximum two passage numbers within the full dataset or within the longitudinal measurements prior to and after BoNT treatment. SCs were differentiated towards myotubes and FI was calculated at two time points with a difference of two passage numbers: no significant alterations of the fusion index at day 6 were reported (n = 5; Supplementary figure 3E).

Characterization of supportive stem cell populations in MG and ST muscles (objective 3.1)

There were no differences between MAB fractions (PDFGRa-ALP+) of TD (MG: 14.27 ± 12.63% and ST: 4.39 ± 4.48%; n = 3) and CP (MG: 15.83 ± 15.16% and ST: 2.27 ± 1.73%; n = 5; Figure 4A). However, a higher MAB proportion was found in the MG muscle compared to ST muscle when pooling data from TD and CP children (n = 8, p = 0.020). FAP fractions (ALP-PDGFRa+) were scarcely represented and showed no significant differences between TD (MG: 0.19 ± 0.12 % and ST: 2.78 ± 4.17%; n = 3) and CP (MG: 0.62 ± 0.69% and ST: 0.87 ± 0.88%; n = 5), nor between muscles (Figure 4B). Fractions of ICs (CD56-ALP-PDGFRa-) were represented with 47.48 ± 5.43% in the MG and 63.37 ± 5.05% in the ST for TD (n = 3) and 36.32 ± 20.86% in the MG and 45.13 ± 23.40% in the ST for CP (n = 5; Figure 4C). ICs representation was higher in the ST muscle compared to MG (n = 8, p = 0.009).

**Figure 4.**
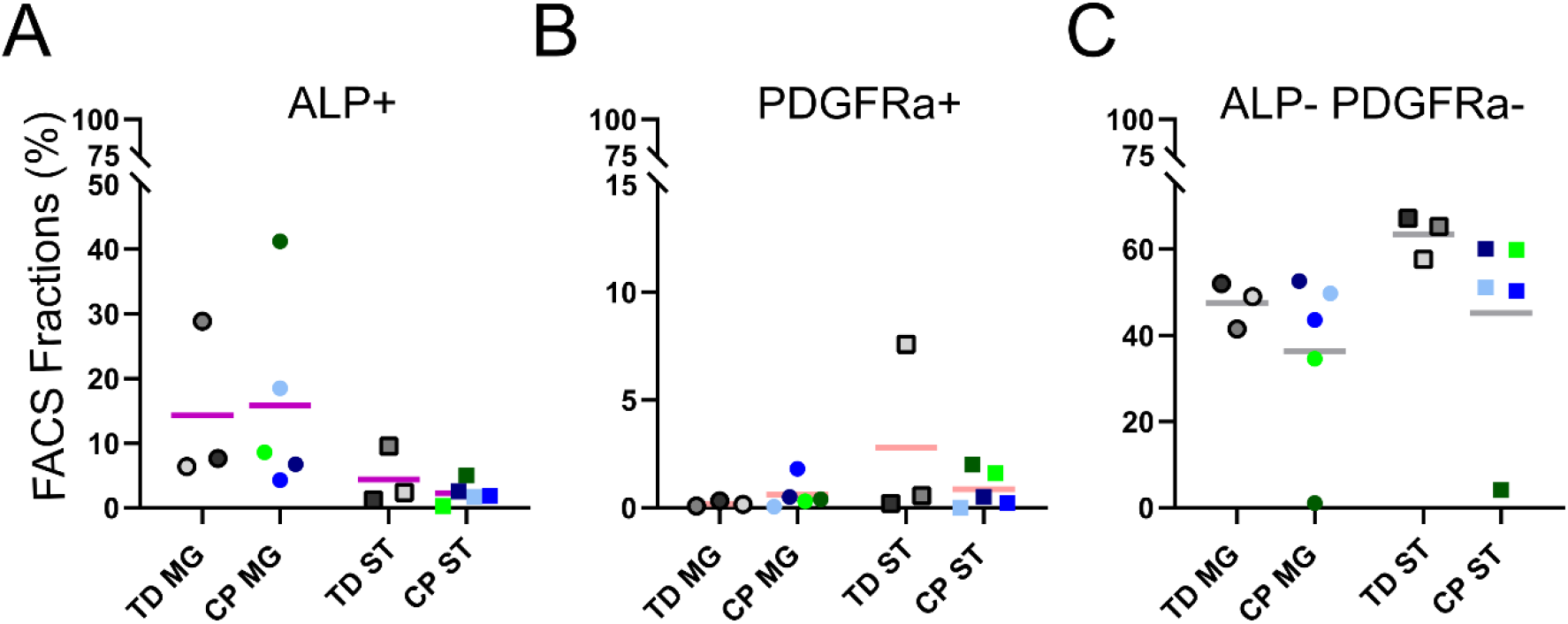
FACS fractions of isolated supportive stem cell populations obtained from the MG and ST muscle in TD and CP children. **(A)** Percentages of obtained CD56-PDGFRa-ALP+ cells based on a serial FACS, referred to as mesoangioblasts (MABs). Data expressed as percentage of total sorted cells. The bar represents the mean. **(B)** Percentages of obtained CD56-ALP-PDGFRa+ cells, referred to as fibro-adipogenic progenitor cells (FAPs), through FACS analyses. **(C)** Percentages of obtained CD56-ALP-PDGFRa-cells, referred to as interstitial cells (ICs), through FACS. Every dot represents data from a different subject, typically developing (TD) children or patients with cerebral palsy (CP) according to the colour code. Cell populations derived from muscle microbiopsies from the MG are indicated by circles, populations from the ST are indicated by squares. The horizontal line represents the mean. TD: n = 3, CP: n = 5

Myogenic differentiation potential of MABs at day 6 of differentiation showed no differences between TD and CP based on MYOD expression and FI values for both muscles (Figure 5A-C). The comparison of MG and ST revealed a significant increase of MYOD+ nuclei in the TD group for the MG (16.44 ± 4.32%) compared to the ST (4.96 ± 2.45%; n = 3; **p < 0.01), while no differences were reported in the CP group (MG: 17.89 ± 16.33%; ST: 10.87 ± 9.26%; n = 5). A high heterogeneity for FI values of the MABs was observed within the CP group and no differences were found between the TD and CP group, nor between both muscles. After adipogenic differentiation of these MABs, similar amounts of lipid content were found for both groups and muscles, while adipogenic capacity slightly increased in the ST muscle of CP compared to TD, but this increase did not reach statistical significance (p = 0.188; Figure 5D, E). FAPs were assessed for adipogenic differentiation potential (Figure 5F, G), as it was previously shown that they do not possess myogenic capacity by themselves ^20^. No differences between TD and CP groups nor between muscles were found, based on ORO quantification. Lastly, ICs (ALP-PDGFRa-) have been assessed for their adipogenic capacity and did not show any significant differences based on ORO quantification between groups nor between muscles (Figure 5H, I). Taken together, we found altered stem cell representations comparing the MG and the ST, with a higher myogenic commitment of the MABs from the MG compared to ST, although no differences in adipogenic potency between muscles nor patient groups were observed.

**Figure 5.**
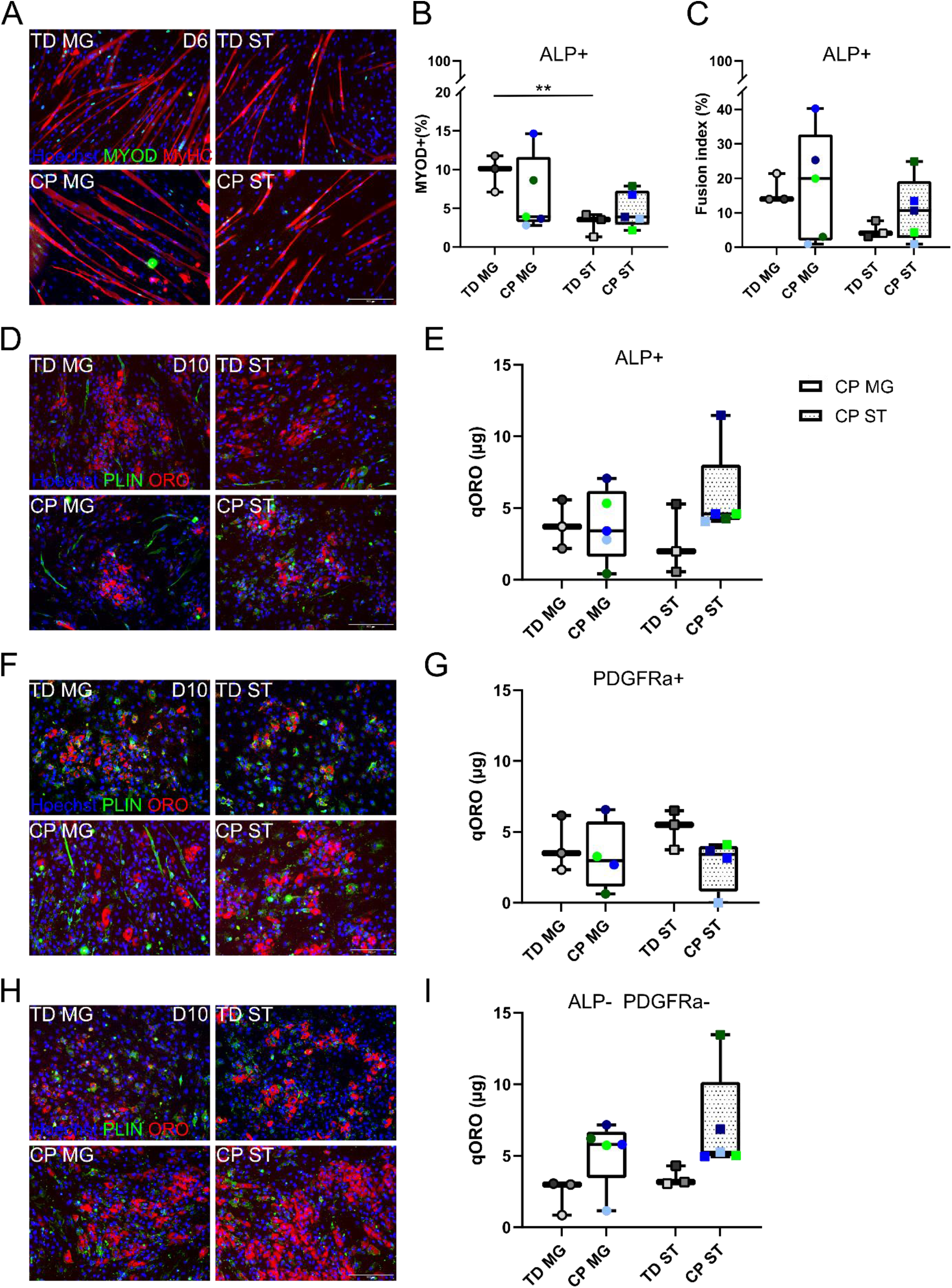
Differentiation potency of MABs, FAPs and ICs from MG and ST muscles of TD children and CP patients. **(A)** Representative IF images from MABs derived from TD children and patients with CP for both MG and ST muscle at day 6 of myogenic differentiation. MYOD (green) and MyHC (red) are shown. Nuclei (blue) are counterstained using Hoechst. **(B)** Percentage of MYOD+ cells based on IF images represented by boxplots for both TD children and CP patients. Dots (MG) and squares (ST) represent individual subjects indicated in different colours. ** p < 0.01 **(C)** MAB (ALP+) fusion indexes shown by boxplots. Dots (MG) and squares (ST) represent individual subjects. **(D)** Representative IF images and **(E)** quantification of ORO based on absorbance at 490nm of MABs from TD children and patients with CP from both MG and ST muscle after adipogenic induction at day 10. PERILIPIN (PLIN, green), Oil red O (ORO, red) are shown. Nuclei (blue) are counterstained using Hoechst. Data expressed in µg of ORO based on a standard curve. **(F)** Representative IF images and **(G)** quantification of ORO of FAPs (PDGFRa+) from TD children and patients with CP from both MG and ST muscle after adipogenic induction at day 10. PLIN (green), ORO (red) and nuclei (blue) are shown. **(H)** Representative IF images and **(I)** quantification of ORO of ICs (ALP-PDGFRa-) from TD children and patients with CP from both MG and ST muscle after adipogenic induction at day 10. PLIN (green), ORO (red) and nuclei (blue) are shown. Scale bars = 200 µm. TD: n = 3, CP: n = 5 (FAP: n = 4)

Of note, for the ST muscle, ICs of one specific patient with CP showed aberrant high adipogenic potency *in vitro* as quantified by ORO. This was linked to the visual assessment of its histological analysis of a muscle section stained with haematoxylin and eosin (Supplementary figure 4). This to qualitatively compare the histological analysis of the ST muscle from a TD child and from another CP child, with average ORO levels for IC adipogenic differentiation. A higher accumulation of fibrosis and/or fatty infiltration by increased connective tissue between the muscle fibers for the sample from this specific CP patient with high IC adipogenic potential, compared to the TD sample and the one CP sample with average ORO values.

### Longitudinal effects on supportive muscle stem cells (objective 3.2)

Examining the effects pre- and post BoNT treatment on the supportive muscle stem cells from CP children showed no differences in FACS fractions for MABs (CD56-ALP+ PDGFRa-cells; n = 4), but revealed a significant decrease for FAPs (CD56-ALP-PDGFRa+ cells; BoNT-naive: 9.13 ± 4.16%; BoNT-treated: 1.04 ± 0.89%; § p < 0.05) and an increase for ICs (CD56-ALP-PDGFRa-cells; BoNT-naive: 10.20 ± 8.62%; BoNT-treated: 42.08 ± 13.65%; § p < 0.05; Figure 6A-C). Based on the repeatability experiments (n = 2), we found that FACS fractions stayed rather consistent for these stem cell populations (average fluctuations between repeated assessments for MABs: 6.44%; FAPs: 0.16% and ICs:14.25%; supplementary figure 5A). To verify whether this small sample size is representative, we portrayed these longitudinal data together with the cross-sectional results of the entire study group. Thereby individual changes between the two timepoints of the subgroup could be compared to the baseline variability amongst all CP children and TD subjects (CP BoNT-naive: n = 6; CP BoNT-treated: n = 9; TD: n = 13; Supplementary figure 5B, C). Additional simple linear regression of these fractions and age was performed on a cross-sectional dataset in an attempt to understand whether these alterations could be due to aging, the progression of CP or associated with BoNT treatment history (Supplementary figure 5D, E). No link with age or BoNT treatment history could be found (n = 11).

**Figure 6.**
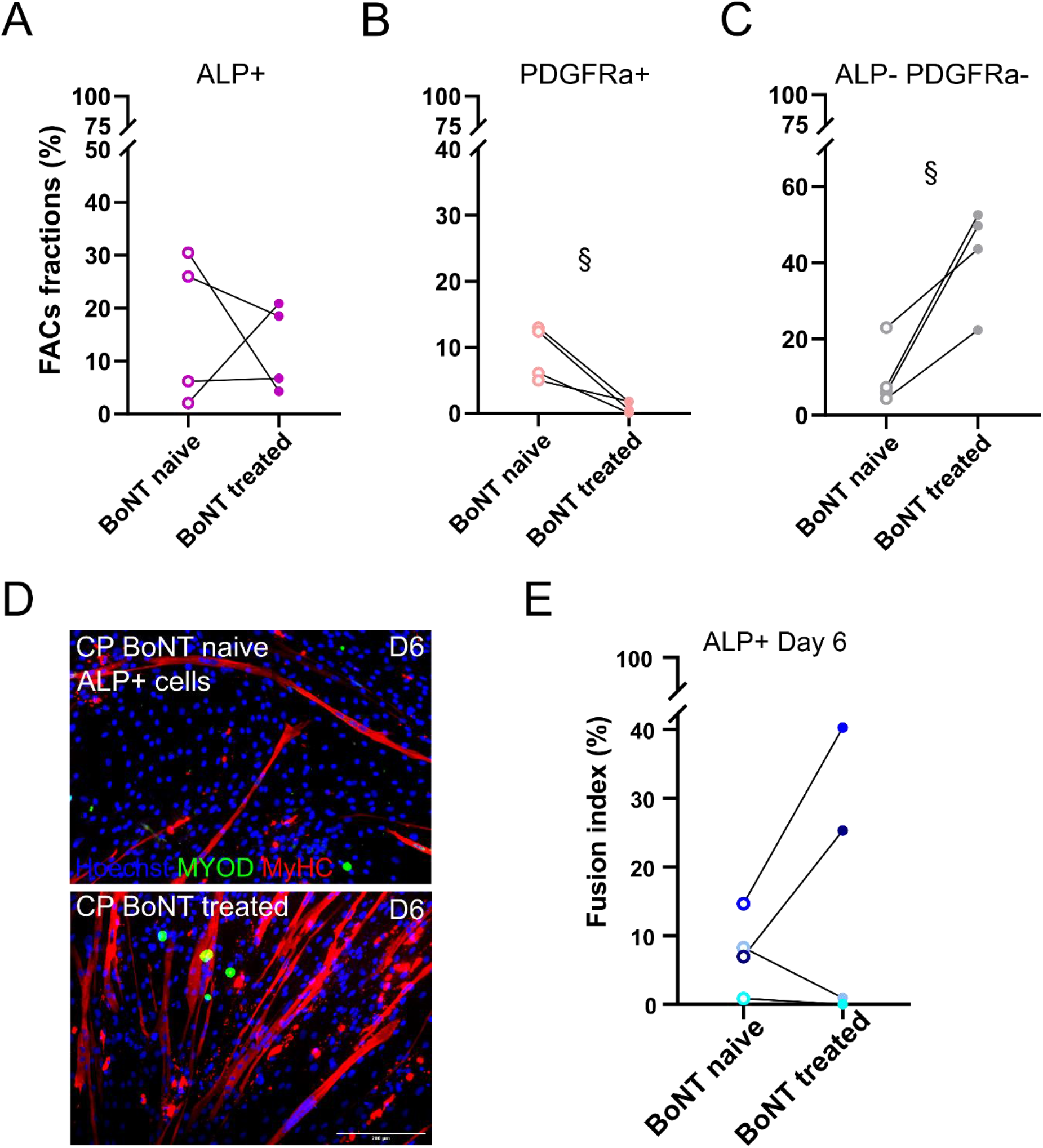
Longitudinal effects on supportive stem cell representations and myogenic capacity in CP children. **(A)** Quantification of serial FACS for MABs (ALP+) from the MG, expressed in percentage of total sorted cells. **(B)** Quantification of fibro-adipogenic progenitor cells (PDGFRa+) from the MG, expressed in percentage of total sorted cells. **(C)** Percentages of obtained ICs (ALP-PDGFRa-). Samples from 4 patients with CP were included before the first Botulinum Neurotoxin A (BoNT) treatment, as well as one year after. Every dot represents an individual subject indicated in different colours. n = 4, § p < 0.05 **(D)** Representative IF images of MABs from patients with CP at day 6 of myogenic differentiation. MYOD (green) and MyHC (red) and nuclei (blue) are shown. Scale bars = 200 µm. **(E)** Fusion indexes of MABs from patients with CP at day 6 of myogenic differentiation. FI of MABs from BoNT naive patients are indicated by dotted lines, after BoNT treatment by solid lines. Colour codes indicate the different subjects (n = 4), all cells were obtained from the MG muscle.

Furthermore, to have a complete overview of myogenic capacity prior to and after BoNT treatment, we assessed MAB myogenic differentiation at day 6. This revealed a steep increase in FI in two out of four cases, but a decrease in fusion potential for the other two cases (Figure 6D, E).

## Discussion

Through the current study three predefined objectives could be achieved. First, we confirmed higher satellite cell (SC) fusion potential from the *Medial Gastrocnemius* (MG) of CP compared to age-matched TD children. Secondly, we reported a decreased myogenic potential of SCs from the *Semitendinosus* (ST) compared to the MG muscle. Furthermore, based on longitudinal assessments, we showed slightly reduced SC representation and fusion capacity, one year after the first BoNT treatment. SC fusion index (FI) values showed to be robust to assess myogenic capacity by repeated analyses. Thirdly, mesoangioblasts (MABs) revealed to be less represented in the ST compared to the MG muscle. Finally, we also described alterations in FACS fractions of supportive stem cell populations after Botulinum Neurotoxin A (BoNT) treatment. These findings will be discussed in detail below, following the structure of objectives provided in table 2.

Increased satellite cell fusion capacity in CP compared to TD (objective 1) Related to the first objective, i.e., comparing SCs from typically developing (TD) children with those of patients with cerebral palsy (CP), we reported higher FI values at day 6 for SCs derived from the MG microbiopsies of young patients with CP compared to age-matched TD children. Thereby, we confirmed our previously published results on SC cultures ^20^. Although very variable results, assessing different muscles and sometimes lacking ideal control data, RNAseq analyses on muscle biopsies have indicated upregulation of pathways related to muscle protein synthesis, indirectly supporting our data on SC differentiation ^39,40^. We have further characterized all cells obtained via the muscle microbiopsy technique that resulted negative for haematopoietic and endothelial cell markers, addressing the concerns about the SC purity recently raised by Lieber and Domenighetti ^21^. These findings were in line with previous reported data, even on additional markers for cells obtained from muscle microbiopsies followed by the explant technique ^41^. Small percentages of cells have been found positive for CD34, which is mainly known as a marker for adult stem cells including a small subset of SCs ^36^. Moreover, cells positive for CD34 have been previously reported after culturing muscle microbiopies ^38,41^. Thereby, we provided evidence that the cells analyzed were not being contaminated by considerable amounts of cells of other lineages.

### Satellite cells features from patients with CP are influenced by multiple variables (objective 2)

To address the second study objective, we assessed multiple factors potentially leading to heterogeneity in reported results on the CP muscle pathology ^19,20,42^. Firstly, to address objective 2.1, we have uniquely cultured and extracted cells from two different muscles of the same young patients with CP, as well as from aged matched TD children. Both, the MG and ST (or entire group of hamstring) muscles have been described at the macroscopic level, i.e. muscle volume via ultrasound assessments and at the microscopic level, i.e. contractile material and collagen content in muscle sections ^4,11,43^, however, rarely from the same subject. Additionally, the majority of previous studies focussed on the entire group of hamstring muscles, without discriminating between the different muscles of the hamstrings ^11,19,32^, as targeting i.e. the ST specifically is more challenging to collect sufficient material due to its smaller size. Thanks to the less invasive character of the muscle microbiopsy technique, we were able to collect and compare muscle samples from young patients with CP, as well as TD children, on stem cells from both muscles, MG and ST, in the same child. This is unique and seems more challenging to achieve with Bergström biopsies or open biopsies during invasive surgery ^4,44^. Here, the focus was primarily on the comparison of data from both muscles, and not anymore on the difference between CP and TD data. Cmparison of both muscles showed a tendency of lowered amounts of SCs (based on FACS fractions) in the ST compared to MG. These tendencies in FACS fractions could be partially elucidated by the activity level of these patients prior to the biopsy ^45,46^. Indeed, physical activity could explain whether these cells were activated and stimulated to self-renew or not as the MG and ST differ in fiber distribution and thereby regeneration potential ^47,48^. Unfortunately, based on their level of physical activities and hobbies, no clear distinctions between the subjects could be made. Additionally, the MG is highly targeted for stretching by physiotherapy and the use of ankle-foot orthoses ^49^, which was indeed also the case in our patient cohort. Possibly, these differences in muscle management between both muscles could have an effect on the observed FACS fractions and other assessed parameters. However, since the slight fluctuations in SC fractions between TD and CP were equally represented in MG compared to ST, we assume that SC *in vitro* amplification was not altered. Possible stimuli from the stem cell niche, or from other cell types, were diminished once the cells were removed from their muscle environment as they were first amplified for 4 passages in culture before sorting, which could have resulted in no significantly altered SC fractions in our study design ^46,50^. Furthermore, studies have reported reduced SC numbers in patients with CP from 10 years of age onwards in different muscles (hamstrings, *Gracilis* and biceps) ^11,15,16^. Yet, no data are available on the age of onset of this phenomenon and on the underlying mechanisms that may potentially exhaust the cells, as is observed in muscular dystrophies ^51^. Additionally, based on IF muscle sections, we could not confirm a lower amount of SCs in CP compared to TD, possibly due to the low sample size and the inclusion of younger subjects. We also did not observe a correlation of *ex vivo* PAX7+ cells with the obtained FACS fractions after *in vitro* amplification. Potentially, the *in vitro* growth conditions could have altered the absolute cell representations as first FACS analyses could only be performed after at least 3 passages, but allow describing rather relative cell representations. Furthermore, it is known that a cell, after extraction, cannot maintain all the molecular and functional properties owned *in vivo* ^52,53^. However, as the expansion method was standardized for all cell lines and included subjects, possible effects on proliferation potential would have led to altered FACS fractions when comparing CP and TD data. Additionally, as mentioned before, even though the cells have been removed from their niche, effects of the *in vivo* situation or of the CP pathology might have influenced their ability to still proliferate *in vitro* ^54^. In this light, even though no direct associations between these *ex vivo* and *in vitro* data could be made, both these data suggest that SC representation was not altered in these patients. The absence of significant alterations supports the hypothesis that, in our cross-sectional study, none of the cell populations within CP muscles were exhausted compared to TD muscles.

We did not find any differences in proliferative capacities based on KI67 expression between TD and CP nor between muscles. KI67 is a commonly used marker for proliferation and is expressed during phases G1, S and G2 of cell cycling, but is absent in resting cells, in G0 ^55,56^. KI67 expression is an indication for the number of cells that are actively involved in cell proliferation, but does not provide information on cell cycling speed, for which more in-depth cell cycle analyses would be required. Previously, another research group showed reduced doubling time for myoblast progenitors derived from hamstring biopsies from patients with CP compared to those of TD adolescents, and higher *KI67* levels based on RNAseq data ^57^. Furthermore, they also stressed the importance of the site for tissue harvesting. They obtained biopsies at the myotendinous junction, which is reported to be richer in proliferating SCs and primarily involved in SC activation and fusion ^58,59^. In contrast, for the current study, muscle microbiopsies were obtained at the muscle mid-belly, contributing to the reported discrepancies between studies ^19,20^. As for myogenic commitment, we found significant higher percentages of MYOD+ nuclei at day 0 of myogenic differentiation in ST SCs compared to MG for TD children, while the opposite trend was found for patients with CP. However, we showed high variance in both MYOD levels and FI values at day 0 between the repeated biopsy samples of two TD children, while these parameters at days 3 and 6 showed to be more robust, letting less conclusive data at day 0. We found significantly lower FI values for the cells extracted from the ST compared to the MG, suggesting lower myogenic potency of these SCs. Interestingly, we measured a FI of around 25% at day 3 of myogenic differentiation of CP-derived SCs from the ST, which is close to the FI value (21%) found by Domenighetti and colleagues after 42h of myogenic differentiation of hamstring myoblasts, regardless many differences in the analysis procedure ^19^. Due to differences in material quantity from the biopsy, Domenighetti group was able to assess myogenic capacity at a younger passage (P4), while we had a longer amplification time and assessed SC differentiation potential at passage 6-8. Differences between the study of Domenighetti group and the current study for the FI values of TD children could potentially be linked to higher age (14 ± 2 years old) and clinical history (ACL reconstructive surgery) and therefore to a potential misuse or altered activity of the muscle. Unfortunately, to our knowledge, no data are available on SCs fusion capacity of healthy subjects of this young age and *in vitro* passage number. Taking the small sample size and heterogeneity of the CP pathology into account, we carefully suggest altered SC features from the MG compared to the ST ^19,20^, hereby highlighting the complexity of CP muscle pathology and rising caution regarding generalized treatment based on muscle-specific findings.

Secondly, although the application of the muscle microbiopsy technique has some previous mentioned limitations, this technique allowed unprecedented follow-up microscopic assessments (to address objective 2.2). Because of the low sample size (partially due to the covid-situation), associations with the BoNT treatment history or the progression of the CP pathology cannot yet be fully explored. Nevertheless, the preliminary results provide some unique first insights in possible alterations over time following the same subject. The number of SCs showed a non-significant decrease one year after BoNT treatment. As was previously shown for SCs of patients within this age range, no correlation with fusion index and age was found ^20^. Although additional data are needed, and potential effects due to activity levels could not be excluded, this preliminary observation is most likely patient-specific and potentially the result of CP progression or the effect of BoNT treatment. Furthermore, myogenic capacity of SCs seemed to have slightly decreased one year after BoNT treatment. This decrease exceeded the repeated assessment variability. As shown, this effect of lower fusion capacity was not associated with the observed lower SC FACS fraction.

Thirdly, to address potential concerns about the assessed outcome parameters, we have performed analyses of two separate muscle biopsies from the same muscle in parallel (objective 2.3). Even though we saw some slight fluctuations in the SC fractions obtained through FACS leading to varying SC yields, the effect on the differences of FI remained low, as we could analyze this parameter within a range of two passages after amplification. Importantly, the differences observed between repeated assessments were lower than the observed differences between TD and CP and between both timepoints during the follow-up, strengthening the fact that the observations we highlight are due to the pathology and not to any other secondary factor.

### Supportive muscle stem cells are not highly altered in patients with CP (objective 3.1)

To address the third study objective, we could successfully obtain multiple muscle stem cell populations next to SCs, based on well-described markers in literature ^20,36^. These results need to be interpreted with caution due to the low sample sizes and the high heterogeneity observed, mainly in the CP group. For both groups, the number of obtained MABs seemed to be lower in the ST compared to the MG muscle, but was not upregulated in CP compared to TD children. This is in contrast to findings on (older) patients with muscular dystrophy disorders ^23,60^. In this regard, we also did not observe a rescue in higher MAB fractions in cases of relatively lower SC numbers ^23,24,60,61^. Further research on larger sample sizes are needed to better understand whether these observations were due to a lack in power or were depending on the age of the enrolled children. However, these data further strengthen the hypothesis of muscle-specific properties and thereby rise caution in interpretation of previous studies considering an entire muscle group or lacking appropriate control data.

The myogenic capacity of MABs, both by MYOD expression and FI, was significantly lower comparing ST-derived cells from TD children and those of the MG. No differences in FI between TD and CP nor between both muscles were found. However, in the same line as what was found for SCs, there was a trend of lower myogenic capacity in the ST for the TD group, while these observations were not confirmed in the CP group, most likely due to higher heterogeneity. Additionally, we did not observe any correlations between the myogenic performance of the SCs and the MABs of the same subjects based on MYOD or FI assessments. Adipogenic potency of these cells did not differ between TD and CP nor between muscles, which was also the case for FAPs and ICs. However, muscle imaging studies on patients with CP have shown higher adipose content in the MG muscle compared to TD subjects in children and adults ^62,63^. Potentially, the subjects in our study were too young to already capture these alterations (*in vitro*), knowing that only a 4% increase in fat fraction was reported in the MG of 8-15-year-olds CP compared to TD ^64^. ICs showed a slight trend of higher adipogenic capacity for the ST muscle compared to the MG, in both groups. This finding may potentially be in line with greater muscle volumes deficits observed in the distal lower limb muscles compared to more proximal ones, as more fat infiltration could compensate potential changes in total muscle volume ^43^. Furthermore, adipogenesis seemed to be linked to a certain location along the muscle, as significant increased fat content was mainly observed at the muscle belly of the MG of more affected patients with CP ^62^. Additionally, in our dataset, one IC cell line from the ST of a patient with CP seems to have a much higher adipogenic potency compared to the other ICs obtained from ST in the CP group that was confirmed on qualitative assessment of histological showing a higher extracellular matrix (ECM) content between the muscle fibers. This assessment included histological imaging of a representative TD and a second patient with CP (matched on GMFCS level and gender). Both patients with CP were not distinguishable based on GMFCS level and strength scores. Interestingly, however based on this single observation, a higher ECM content was observed for the patient with repetitive BoNT treatments and slightly older in age (22 months) compared to the other patient with CP, which only received one previous BoNT treatment. More quantitative data as well as more specific staining for fat infiltrations would be required to give robust associations to our described single case. More specific investigations will be able to give a better interpretation of these observations, seen the importance of understanding fat infiltration mechanisms together with progression of this as many other diseases affecting skeletal muscle.

### Longitudinal effects on supportive stem cells pre- and post BoNT in patients with CP (objective 3.2)

Based on the FACS data of all four patients, we showed a significant decrease in FAP fraction and an increase in IC fraction in CP children after one year. It has been shown that, in chronic atrophic conditions caused by motor neuron deficits, increased fibrosis is associated with accumulation of FAPs in the interstitium of denervated muscles ^65– 67^. This seemed not to be the case by our FACS fractions, even though the CP pathology is expected to be characterized by increased fibrotic content ^68^. The included subjects may have been too young to already notice differences in active involvement of these stem cell populations as we currently do not know at which age this accumulation in ECM occurs in the muscle and if it occurs differently from muscle to muscle. Additionally, as ICs are defined by negative expression for markers CD56, ALP and PDGFRa, possible intrinsic differences in this mixed population remained unnoticed. The reduction in PDGFRa population (FAPs) could have contributed to the increase of the IC population, potentially incrementing ECM deposition ^69^. Moreover, alterations in PDGF signaling could modulate multiple processes in cell survival, cell fate decisions and damage-associated behaviours in the muscle, contributing to the observed CP muscle pathology ^70^.

## Conclusion

This study confirmed our previous findings that satellite cell (SC) cultures derived from the *Medial Gastrocnemius* (MG) muscle of young patients with cerebral palsy (CP) have a higher fusion capacity compared to those of age-matched peers. Through enlarging the sample size, clarifying the sample identity and refining the sample homogeneity, we obtained results that are in line with what was previously published ^20^. Parameters were proven to be quite robust based on repeatability experiments assessing SCs in parallel from the same subject and muscle. As previously underlined, we are sharing the idea that unravelling the effects of multiple factors, such as assessed muscle, progression and reliability of the primary outcome parameters, will help to understand the observed heterogeneity raised between our and previously published reports. Therefore, for the first time, obtaining cells through the muscle microbiopsy technique from two different lower limb muscles of the same subject, allowed a better improvement in the knowledge of the muscle heterogeneity. Myogenic potential of both SCs and mesoangioblasts seemed to be decreased in the *Semitendinosus* (ST) compared to the MG muscle, rising caution to draw general conclusions and develop novel general treatments ^57^. The low sample size and the limited possibility to compare our results with few previously published data still need to be improved. Nevertheless, our data are sharpening our efforts towards understanding the differences in proliferation and differentiation abilities of muscle cells progressing together with CP pathology. Indeed, unprecedented preliminary longitudinal data showed altered stem cell population proportions, as well as slight reduction in SC myogenic capacity one year after the first treatment with Botulinum Neurotoxin. Future research will be focussing on the observed stem cell phenotype, muscle composition and in combination with additional clinical data, this may lead to a further understanding of the development and complexity of CP muscle pathology. Ultimately, in the spirit to work on our experiments and obtain more data uncovering the insights into CP pathology, we offer one step further to pave the way for more optimal treatment approaches for patients with CP.

## Declarations

### Ethics approval and consent to participate

The studies involving human participants were reviewed and approved by the Ethical Committee of the University Hospitals of Leuven, Belgium (S61110 and S62645). Written informed consent to participate in this study was provided by the participants’ legal guardian/next of kin.

### Authors’ contributions

All authors conceived and discussed experiments, read and approved the final version of the manuscript. In particular, EDW and AVC are pediatric orthopedic surgeons, collecting the muscle microbiopsies and together with EH were responsible for patient recruitment. MC, JM, JD and DC performed the experiments. MC, JM, and DC analyzed the data and prepared the figures. MC wrote the manuscript. GG, MS, AVC, KD and DC edited and revised the manuscript.

## Acknowledgements

We warmly thank all children and their families for participating in this study. We are grateful for the contribution of Prof. MD S. Nijs and E. Nijs from the Traumatology Department, Prof. G. Molenaers and Dr. H. De Houwer from the paediatric orthopaedic department and Prof. MD G. Hens from the Ear, Nose Throat department for the contribution to the recruitment of TD children, associated to the University Hospital of Leuven, Belgium, as well as all staff involved. Special thanks to J. Uytterhoeven and L. Staut, who supported the recruitment procedure. We also want to express our gratitude towards S. Vlayen and R. Ramezankhani for providing the cell lines as positive controls for the FACS analyses.

## References

1. Oskoui, M., Coutinho, F., Dykeman, J., Jetté, N. & Pringsheim, T. An update on the prevalence of cerebral palsy: a systematic review and meta-analysis. Dev. Med. Child Neurol. 55, 509–519 (2013).

2. Mathewson, M. A. & Lieber, R. L. Pathophysiology of Muscle Contractures in Cerebral Palsy. Phys. Med. Rehabil. Clin. N. Am. 26, 57–67 (2015).

3. Romero, B., Robinson, K. G., Batish, M. & Akins, R. E. An emerging role for epigenetics in cerebral palsy. J. Pers. Med. 11, (2021).

4. Howard, J. J., Graham, K. & Shortland, A. P. Understanding skeletal muscle in cerebral palsy: a path to personalized medicine? Dev. Med. Child Neurol. 64, 289–295 (2022).

5. Palisano, R. et al. Development and reliability of a system to classify gross motor function in children with cerebral palsy. Dev. Med. Child Neurol. 39, 214– 223 (2008).

6. Palisano, R. J., Rosenbaum, P., Bartlett, D. & Livingston, M. H. Content validity of the expanded and revised Gross Motor Function Classification System. Dev. Med. Child Neurol. 50, 744–750 (2008).

7. Novak, I. Evidence-Based Diagnosis, Health Care, and Rehabilitation for Children With Cerebral Palsy. J. Child Neurol. 29, 1141–1156 (2014).

8. Smith, L. R. et al. Contribution of extracellular matrix components to the stiffness of skeletal muscle contractures in patients with cerebral palsy. Connect. Tissue Res. (2019). doi:10.1080/03008207.2019.1694011

9. Pingel, J. et al. Gene expressions in cerebral palsy subjects reveal structural and functional changes in the gastrocnemius muscle that are closely associated with passive muscle stiffness. Cell Tissue Res. 384, 513–526 (2021).

10. Booth, C. M., Cortina-Borja, M. J. & Theologis, T. N. Collagen accumulation in muscles of children with cerebral palsy and correlation with severity of spasticity. Dev. Med. Child Neurol. 43, 314–20 (2001).

11. Smith, L. R., Lee, K. S., Ward, S. R., Chambers, H. G. & Lieber, R. L. Hamstring contractures in children with spastic cerebral palsy result from a stiffer extracellular matrix and increased in vivo sarcomere length. J. Physiol. 589, 2625–39 (2011).

12. Marbini, A. et al. Immunohistochemical study of muscle biopsy in children with cerebral palsy. Brain Dev. 24, 63–66 (2002).

13. Johnson, D. L., Miller, F., Subramanian, P. & Modlesky, C. M. Adipose tissue infiltration of skeletal muscle in children with cerebral palsy. J. Pediatr. 154, 715–720 (2009).

14. Smith, L. R., Chambers, H. G. & Lieber, R. L. Reduced satellite cell population may lead to contractures in children with cerebral palsy. Dev. Med. Child Neurol. 55, 264–270 (2013).

15. Dayanidhi, S. et al. Reduced satellite cell number in situ in muscular contractures from children with cerebral palsy. J. Orthop. Res. 33, 1039–1045 (2015).

16. Von Walden, F. et al. Muscle contractures in patients with cerebral palsy and acquired brain injury are associated with extracellular matrix expansion, pro-inflammatory gene expression, and reduced rRNA synthesis. Muscle and Nerve 58, 277–285 (2018).

17. Fu, X., Wang, H. & Hu, P. Stem cell activation in skeletal muscle regeneration. Cellular and Molecular Life Sciences 72, 1663–1677 (2015).

18. Bachman, J. F. & Chakkalakal, J. V. Insights into muscle stem cell dynamics during postnatal development. FEBS J. 1–13 (2021). doi:10.1111/febs.15856

19. Domenighetti, A. A. et al. Loss of myogenic potential and fusion capacity of muscle stem cells isolated from contractured muscle in children with cerebral palsy. Am J Physiol Cell Physiol 315, 247–257 (2018).

20. Corvelyn, M. et al. Muscle Microbiopsy to Delineate Stem Cell Involvement in Young Patients: A Novel Approach for Children With Cerebral Palsy. Front. Physiol. 11, (2020).

21. Lieber, R. L. & Domenighetti, A. A. Commentary: Muscle Microbiopsy to Delineate Stem Cell Involvement in Young Patients: A Novel Approach for Children With Cerebral Palsy. Front. Physiol. 12, 945 (2021).

22. Sampaolesi, M. et al. Mesoangioblast stem cells ameliorate muscle function in dystrophic dogs. Nature 444, 574–579 (2006).

23. Dellavalle, A. et al. Pericytes resident in postnatal skeletal muscle differentiate into muscle fibres and generate satellite cells. Nat. Commun. 2, 499 (2011).

24. Tedesco, F. S., Moyle, L. A. & Perdiguero, E. Muscle Interstitial Cells: A Brief Field Guide to Non-satellite Cell Populations in Skeletal Muscle. in Methods in molecular biology (Clifton, N.J.) 1556, 129–147 (2017).

25. Joe, A. W. B. et al. Muscle injury activates resident fibro/adipogenic progenitors that facilitate myogenesis. Nat. Cell Biol. 12, 153–163 (2010).

26. Uezumi, A., Fukada, S., Yamamoto, N., Takeda, S. & Tsuchida, K. Mesenchymal progenitors distinct from satellite cells contribute to ectopic fat cell formation in skeletal muscle. Nat. Cell Biol. 12, 143–152 (2010).

27. Collins, B. C. & Kardon, G. It takes all kinds: heterogeneity among satellite cells and fibro-adipogenic progenitors during skeletal muscle regeneration. Development 148, (2021).

28. Hogarth, M. W. et al. Fibroadipogenic progenitors are responsible for muscle loss in limb girdle muscular dystrophy 2B. Nat. Commun. 10, 2430 (2019).

29. De Bruin, M., Smeulders, M. J., Kreulen, M., Huijing, P. A. & Jaspers, R. T. Intramuscular connective tissue differences in spastic and control muscle: A mechanical and histological study. PLoS One 9, (2014).

30. Lieber, R. L. & Fridén, J. Muscle contracture and passive mechanics in cerebral palsy. J. Appl. Physiol. 126, 1492–1501 (2019).

31. Hösl, M. et al. Impact of Altered Gastrocnemius Morphometrics and Fascicle Behavior on Walking Patterns in Children With Spastic Cerebral Palsy. Front. Physiol. 11, 518134 (2020).

32. Htwe, O. et al. Urine hydroxyproline correlates with progression of spasticity in cerebral palsy. Electron. J. Gen. Med. 15, 1–9 (2017).

33. Walhain, F., Desloovere, K., Declerck, M., Van Campenhout, A. & Bar-On, L. Interventions and lower-limb macroscopic muscle morphology in children with spastic cerebral palsy: a scoping review. Dev. Med. Child Neurol. 63, 274–286 (2021).

34. Sätilä, H. Over 25 Years of Pediatric Botulinum Toxin Treatments: What Have We Learned from Injection Techniques, Doses, Dilutions, and Recovery of Repeated Injections? Toxins (Basel). 12, 440 (2020).

35. Kinney, M. C. et al. Reduced skeletal muscle satellite cell number alters muscle morphology after chronic stretch but allows limited serial sarcomere addition. Muscle Nerve 55, 384–392 (2017).

36. Tey, S. R., Robertson, S., Lynch, E. & Suzuki, M. Coding Cell Identity of Human Skeletal Muscle Progenitor Cells Using Cell Surface Markers: Current Status and Remaining Challenges for Characterization and Isolation. Frontiers in Cell and Developmental Biology 7, (2019).

37. Zhou, Q., Yao, Y. & Ericson, S. G. The Protein Tyrosine Phosphatase CD45 Is Required for Interleukin 6 Signaling in U266 Myeloma Cells. Int. J. Hematol. 79, 63–73 (2004).

38. Sidney, L. E., Branch, M. J., Dunphy, S. E., Dua, H. S. & Hopkinson, A. Concise Review: Evidence for CD34 as a Common Marker for Diverse Progenitors. Stem Cells 32, 1380–1389 (2014).

39. Smith, L. R., Chambers, H. G., Subramaniam, S. & Lieber, R. L. Transcriptional abnormalities of hamstring muscle contractures in children with cerebral palsy. PLoS One 7, (2012).

40. Robinson, K. G., Crowgey, E. L., Lee, S. K. & Akins, R. E. Transcriptional analysis of muscle tissue and isolated satellite cells in spastic cerebral palsy. Dev. Med. Child Neurol. 63, 1213–1220 (2021).

41. Ceusters, J. et al. From skeletal muscle to stem cells: An innovative and minimally-invasive process for multiple species. Sci. Rep. 7, (2017).

42. Handsfield, G. G., Williams, S., Khuu, S., Lichtwark, G. & Stott, N. S. Muscle architecture, growth, and biological Remodelling in cerebral palsy: a narrative review. BMC Musculoskelet. Disord. 23, 233 (2022).

43. Modlesky, C. M. & Zhang, C. Muscle Size, Composition, and Architecture in Cerebral Palsy. in Cerebral Palsy 253–268 (Springer International Publishing, 2020). doi:10.1007/978-3-319-74558-9_14

44. Bergström, J. & Edwards, R. H. T. MUSCLE-BIOPSY NEEDLES. The Lancet 313, 153 (1979).

45. Snijders, T. et al. Satellite cells in human skeletal muscle plasticity. Front. Physiol. 6, 1–21 (2015).

46. Chen, W., Datzkiw, D. & Rudnicki, M. A. Satellite cells in ageing: use it or lose it. Open Biol. 10, 200048 (2020).

47. Zimowska, M. et al. Inflammatory response during slow- and fast-twitch muscle regeneration. Muscle Nerve 55, 400–409 (2017).

48. Sharma, G. R., Kumar, V., Kanojia, R. K., Vaiphei, K. & Kansal, R. Fast and slow myosin as markers of muscle regeneration in mangled extremities: a pilot study. Eur. J. Orthop. Surg. Traumatol. 29, 1539–1547 (2019).

49. Hösl, M., Böhm, H., Arampatzis, A. & Döderlein, L. Effects of ankle – foot braces on medial gastrocnemius morphometrics and gait in children with cerebral palsy. J. Child. Orthop. 209–219 (2015). doi:10.1007/s11832-015-0664-x

50. Cornelison, D. D. W. Context matters: In vivo and in vitro influences on muscle satellite cell activity. J. Cell. Biochem. 105, 663–669 (2008).

51. Chang, N. C., Chevalier, F. P. & Rudnicki, M. A. Satellite Cells in Muscular Dystrophy – Lost in Polarity. Trends Mol. Med. 22, 479–496 (2016).

52. Sharifiaghdas, F., Taheri, M. & Moghadasali, R. Isolation of human adult stem cells from muscle biopsy for future treatment of urinary incontinence. Urol. J. 8, 54–9 (2011).

53. Saccone, V. et al. HDAC-regulated myomiRs control BAF60 variant exchange and direct the functional phenotype of fibro-adipogenic progenitors in dystrophic muscles. Genes Dev. 28, 841–857 (2014).

54. Yin, H., Price, F. & Rudnicki, M. A. Satellite Cells and the Muscle Stem Cell Niche. Physiol. Rev. 93, 23–67 (2013).

55. Mackey, A. L. et al. Assessment of satellite cell number and activity status in human skeletal muscle biopsies. Muscle Nerve 40, 455–465 (2009).

56. Tan, C. M. et al. Modulation of Ki67 and myogenic regulatory factor expression by tocotrienol-rich fraction ameliorates myogenic program of senescent human myoblasts. Arch. Med. Sci. 17, 752–763 (2021).

57. Sibley, L. A. et al. Differential DNA methylation and transcriptional signatures characterize impairment of muscle stem cells in pediatric human muscle contractures after brain injury. FASEB J. 35, 1–17 (2021).

58. Allouh, M. Z., Yablonka-Reuveni, Z. & Rosser, B. W. C. Pax7 Reveals a Greater Frequency and Concentration of Satellite Cells at the Ends of Growing Skeletal Muscle Fibers. J. Histochem. Cytochem. 56, 77–87 (2008).

59. Kim, M. et al. Single-nucleus transcriptomics reveals functional compartmentalization in syncytial skeletal muscle cells. Nat. Commun. 11, 6375 (2020).

60. Díaz-Manera, J. et al. The increase of pericyte population in human neuromuscular disorders supports their role in muscle regeneration in vivo. J. Pathol. 228, 544–553 (2012).

61. Kuang, S., Gillespie, M. A. & Rudnicki, M. A. Niche Regulation of Muscle Satellite Cell Self-Renewal and Differentiation. Cell Stem Cell 2, 22–31 (2008).

62. D’Souza, A., Bolsterlee, B., Lancaster, A. & Herbert, R. D. Intramuscular fat in children with unilateral cerebral palsy. Clin. Biomech. 80, 105183 (2020).

63. Svane, C. et al. Quantitative MRI and Clinical Assessment of Muscle Function in Adults With Cerebral Palsy. Front. Neurol. 12, 771375 (2021).

64. Akinci D’Antonoli, T. et al. Combination of Quantitative MRI Fat Fraction and Texture Analysis to Evaluate Spastic Muscles of Children With Cerebral Palsy. Front. Neurol. 12, 633808 (2021).

65. Contreras, O., Rebolledo, D. L., Oyarzún, J. E., Olguín, H. C. & Brandan, E. Connective tissue cells expressing fibro/adipogenic progenitor markers increase under chronic damage: relevance in fibroblast-myofibroblast differentiation and skeletal muscle fibrosis. Cell Tissue Res. 364, 647–660 (2016).

66. Madaro, L. et al. Denervation-activated STAT3–IL-6 signalling in fibro-adipogenic progenitors promotes myofibres atrophy and fibrosis. Nat. Cell Biol. 20, 917–927 (2018).

67. Biferali, B., Proietti, D., Mozzetta, C. & Madaro, L. Fibro–Adipogenic Progenitors Cross-Talk in Skeletal Muscle: The Social Network. Front. Physiol. 10, (2019).

68. Howard, J. J. & Herzog, W. Skeletal Muscle in Cerebral Palsy: From Belly to Myofibril. Front. Neurol. 12, (2021).

69. Shadrin, I. Y., Khodabukus, A. & Bursac, N. Striated muscle function, regeneration, and repair. Cell. Mol. Life Sci. 73, 4175–4202 (2016).

70. Contreras, O., Rossi, F. M. V & Theret, M. Origins, potency, and heterogeneity of skeletal muscle fibro-adipogenic progenitors—time for new definitions. Skelet. Muscle 11, 16 (2021).

