## Supplementary Table 1 for "Adult stem cell characterization from the *Medial Gastrocnemius* and *Semitendinosus* muscles in early development of cerebral palsy pathology"

**Supplementary Table 1. List of used antibodies.**

| **Antibody** | **Relative dilution** | **Provider** | **#Catalog number clone** | **Application** |
|---|---|---|---|---|
| CD56-APC | 0.1 µL/1.$10^6$ cells | Biolegend | #304610 MEM-188 | FACS |
| ALP-PE | 1.2 µL/1.$10^6$ cells | R&D Systems | #FAB1448P B4-78 | FACS |
| PDGFRa-APC | 0.1 µL/1.$10^6$ cells | Biolegend | #323512 16A1 | FACS |
| CD34-APC | 0.5 µL/3.$10^5$ cells | eBioscience | #17-0349-42 4H11 | Flow cytometry |
| CD45-APC | 0.5 µL/3.$10^5$ cells | eBioscience | #17-9459-42 2D1 | Flow cytometry |
| CD144-APC | 0.5 µL/3.$10^5$ cells | Biolegend | #348508 BV9 | Flow cytometry |
| CD31-PE | 0.1 µL/3.$10^5$ cells | Biolegend | #303106 WM59 | Flow cytometry |
| MyHC | 1:20 | Hybridoma Bank | MF20 (mouse) | IF staining (cell culture) |
| MYOD | 1:200 | Cell Signaling | #13812S D8G3 (rabbit) | IF staining (cell culture) |
| KI67 | 1:300 | BD Pharmingen | #556003 B56 (mouse) | IF staining (cell culture) |
| PLIN | 1:200 | Merck | #P1873 polyclonal (rabbit) | IF staining (cell culture) |
| PAX7 | 1:2 | Hybridoma Bank | DSHB Pax7 (mouse) | IF staining (muscle section) |
| LAMININ | 1:250 | Abcam | #ab11575 polyclonal (rabbit) | IF staining (muscle section) |

*ALP: ALKALINE PHOSPHATASE, PDGFRa: PLATELET-DERIVED GROWTH FACTOR RECEPTOR alpha, MyHC: MYOSIN HEAVY CHAIN, PLIN: PERILIPIN, FACS: Fluorescence-activated cell sorting, IF: Immunofluorescent*
