## Supplementary figure 1 for "Adult stem cell characterization from the *Medial Gastrocnemius* and *Semitendinosus* muscles in early development of cerebral palsy pathology"

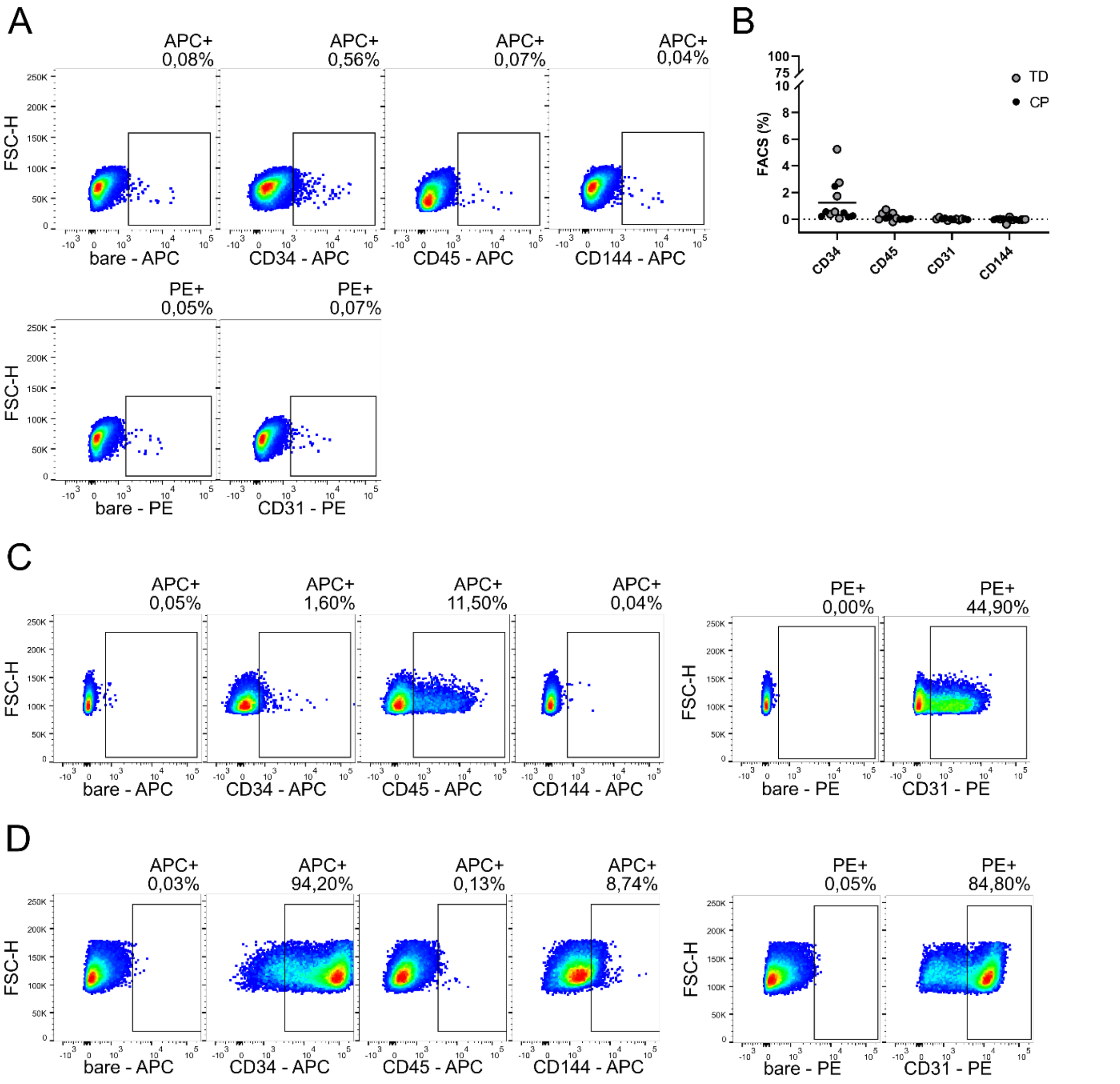


**Supplementary Figure 1. FACS characterization of cultured muscle cells. (A)** Representative example of Fluorescent activated cell sorting (FACS) characterization of all cultured cells derived from muscle microbiopsies of the *Medial Gastrocnemius*. Both left graphs show the unstained sample for the APC- and PE-channel, respectively. Analysis was performed using markers CD34, CD45, CD144 and CD31. **(B)** Percentages of positive populations are represented in the graph. Every dot refers to a sample from a typically developing child (TD; n = 6), or patient with cerebral palsy (CP; n = 6). One sample T-tests were performed for significant differences from 0. CD34 had a p-value of 0.0193 and was the only significant value. **(C)** FACS analysis of positive control sample consisting of lymphocytes from the U266 cell line analyzed for CD34, CD45, CD144 and CD31. Both graphs gating for channels APC and PE on unstained cells are shown. Percentages are indicated on the graphs. **(D)** FACS analysis for positive control sample consisting of human induced pluripotent stem cells (hiPSCs)-derived endothelial cells analyzed for CD34, CD45, CD144 and CD31. Both graphs gating for channels APC and PE on unstained cells are shown.
