## Supplementary figure 2 for "Adult stem cell characterization from the *Medial Gastrocnemius* and *Semitendinosus* muscles in early development of cerebral palsy pathology"

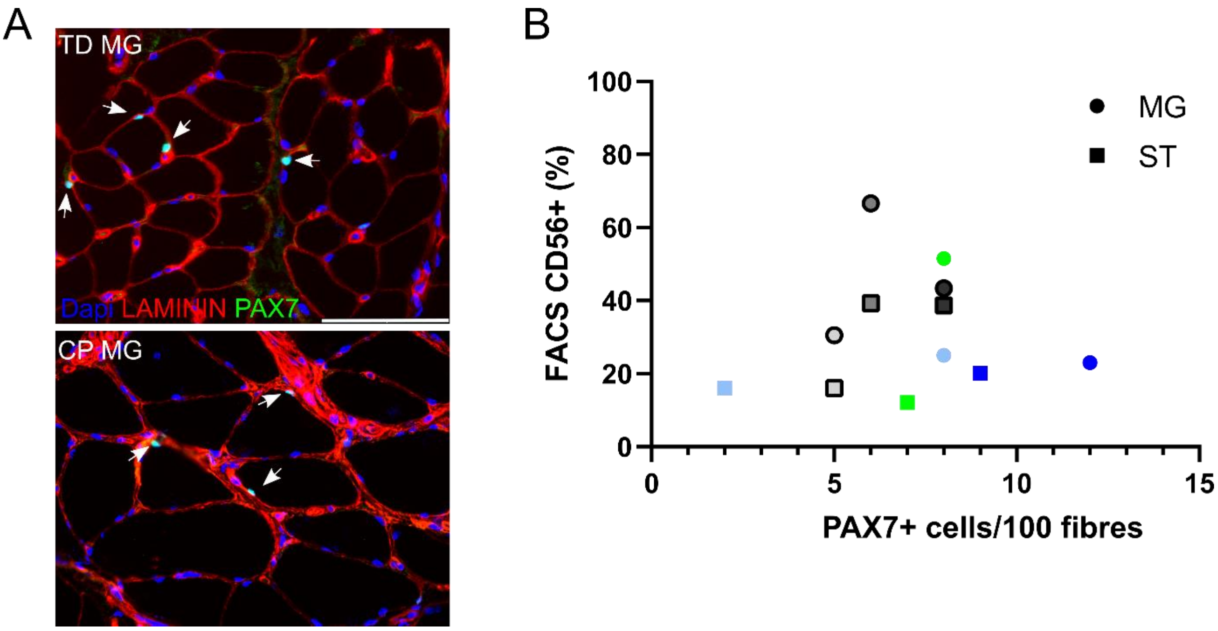


**Supplementary Figure 2. FACS correlation to *ex vivo* presence of satellite cells. (A)** Representative immunofluorescent (IF) images of MG muscle sections from both TD children and patients with CP. LAMININ (red) and PAX7 (green) are shown. Nuclei (blue) are counterstained using Dapi. Arrows indicate SCs, by co-localization of PAX7 and Dapi. Scale bar = 100 µm. **(B)** Correlation of SCs obtained through FACS (CD56+) and the number of PAX7+ cells counted via IF in muscle sections, normalized per 100 fibers. Data of the MG are indicated by dots, data of ST muscle by squares. Gray shades indicate TD data, colours have been used for CP data, following the colour code (n = 12).
