## Supplementary figure 3 for "Adult stem cell characterization from the *Medial Gastrocnemius* and *Semitendinosus* muscles in early development of cerebral palsy pathology"

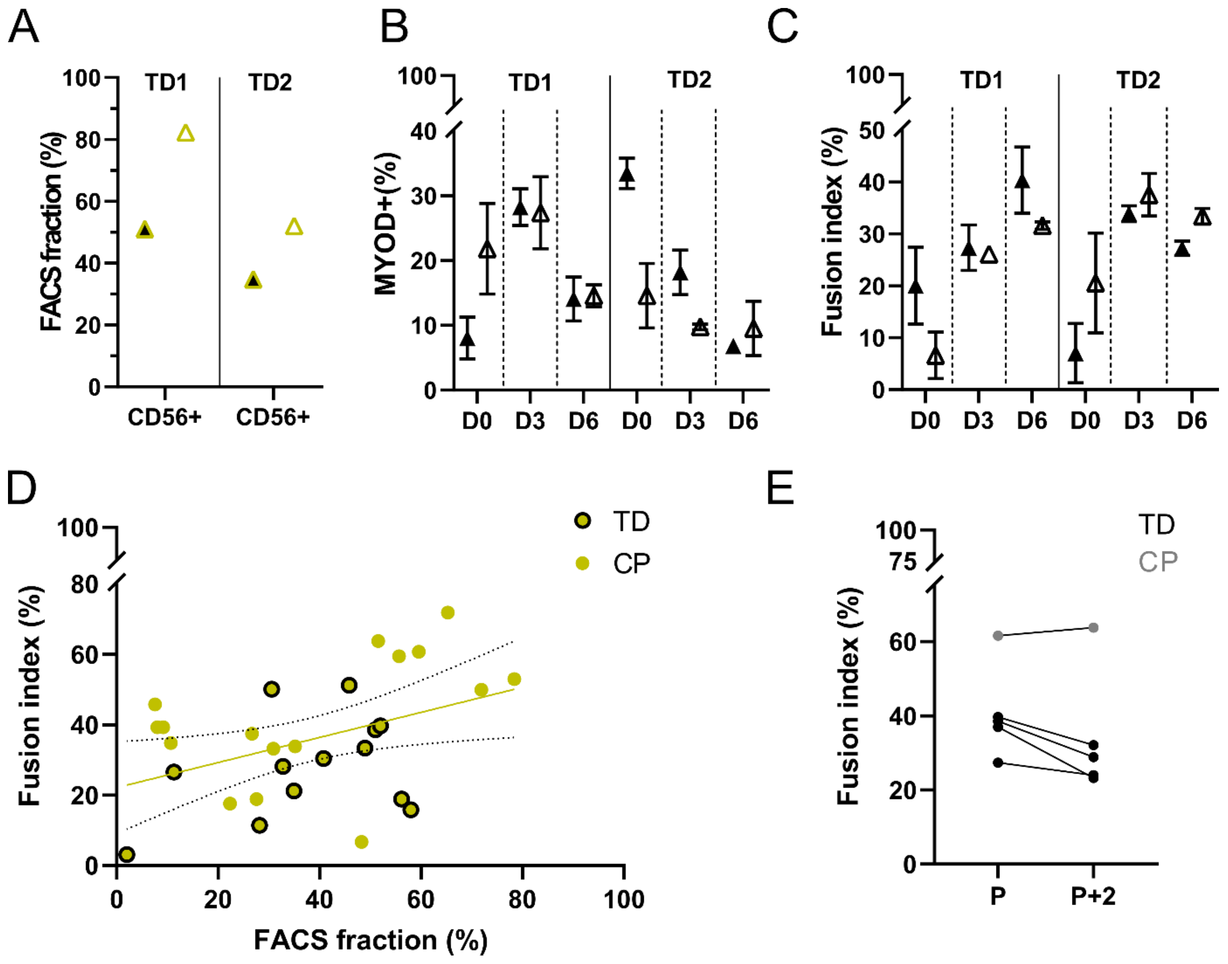


**Supplementary figure 3. Procedural effects on SC characteristics. (A)** Repeated assessments of CD56+, SCs, fractions from two separate microbiopsies obtained in parallel from the same subject (TD n = 2). **(B)** Repeated assessments for percentage of MYOD+ cells of SCs during myogenic differentiation at days 0, 3 and 6, based on 3 representative IF images. SCs were obtained from the same subject on parallel microbiopsies of the MG. **(C)** Repeated assessments for fusion indexes of SCs during myogenic differentiation at days 0, 3 and 6, based on 3 representative IF images. **(D)** Linear regression for SC fusion index at day 6 and FACS fractions for CD56+ cells. Trend line and 95% confidence interval are indicated. Dots with black border represent individual TD subjects, while closed dots represent patients with CP. $R^2$: 0,1781 (TD n = 13, CP n = 16). **(E)** Fusion indexes from the same CD56+ cell lines at day 6 of myogenic differentiation with a time interval of 2 passages (P). TD-derived data are represented by black dots, CP by gray dots, the same subjects are connected by a black line (n = 5).
