## Supplementary figure 4 for "Adult stem cell characterization from the *Medial Gastrocnemius* and *Semitendinosus* muscles in early development of cerebral palsy pathology"

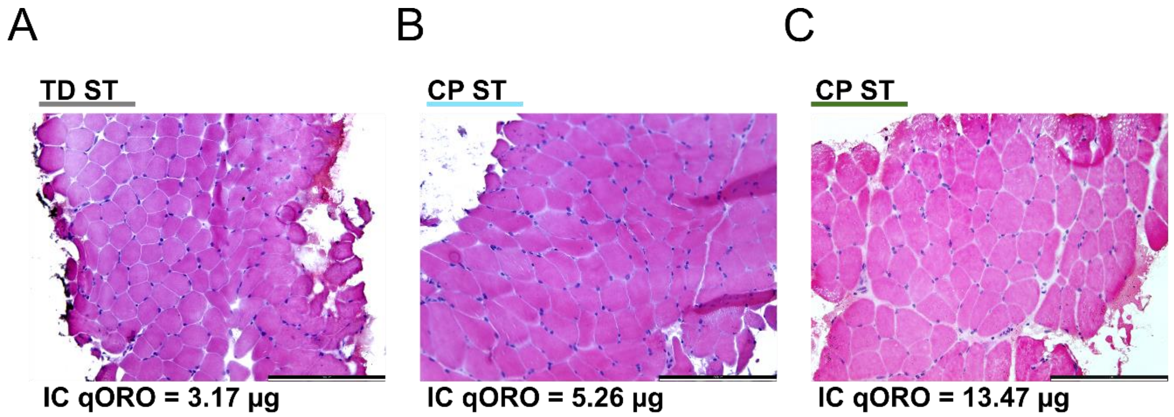


**Supplementary figure 4.** ***Ex vivo*** **representative imaging of muscle integrity from a TD child and two patients with CP. (A)**. Haematoxylin and eosin (H&E) staining from ST muscle slice from a representative TD (colour code is used for underlining). Quantification of Oil Red O (qORO) staining after *in vitro* adipogenic differentiation of interstitial cell (IC) population is shown. **(B)** H&E staining of ST muscle from patient with CP with average ORO levels. **(C)** H&E staining from ST muscle of CP patient with high adipogenic potency for ICs (figure 3I). Scale bars = 200 µm.
