## Supplementary figure 5 for "Adult stem cell characterization from the *Medial Gastrocnemius* and *Semitendinosus* muscles in early development of cerebral palsy pathology"

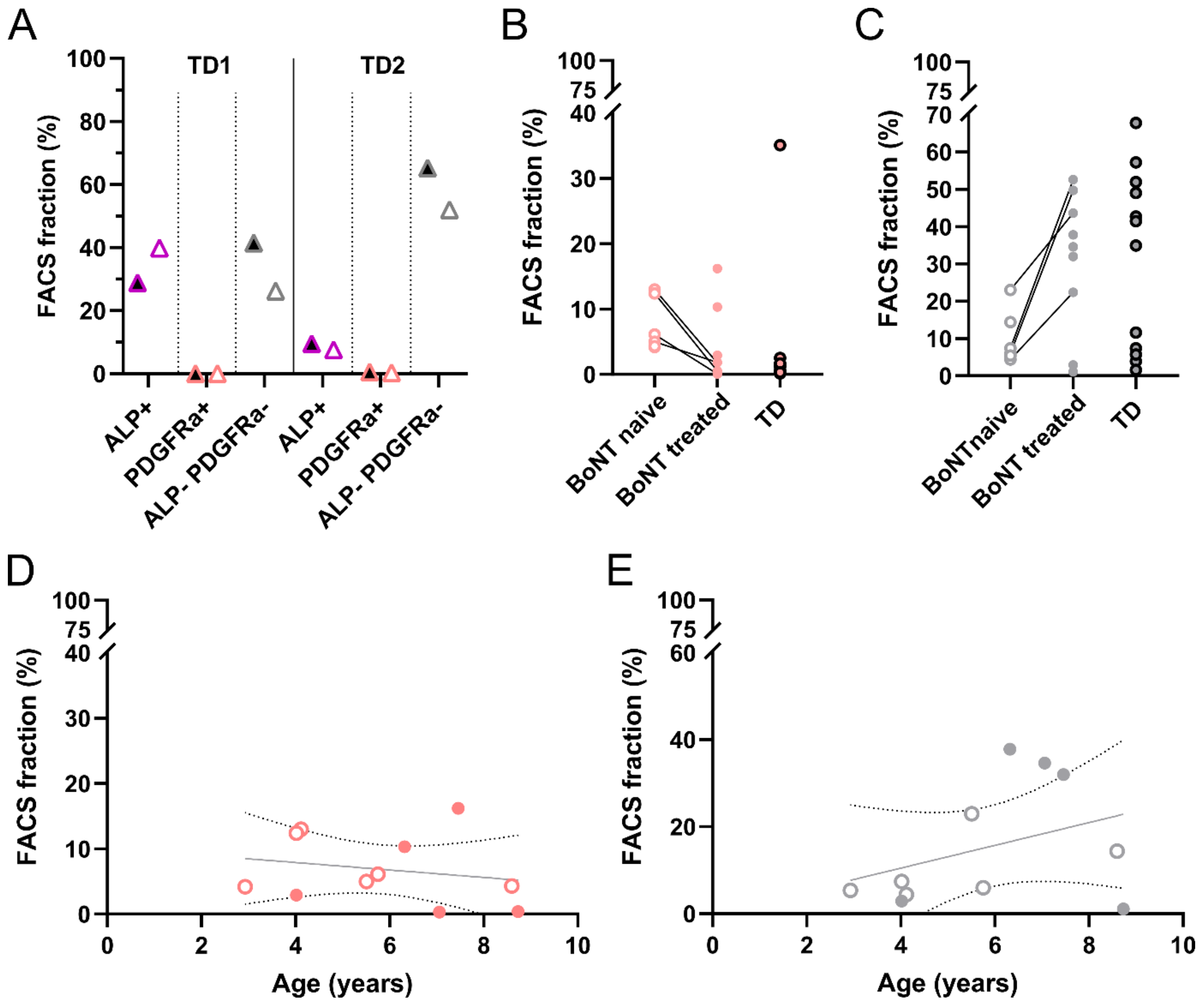


**Supplementary figure 5. Cross-sectional FACS data of supportive stem cell populations to frame obtained results. (A)** Repeated assessments of MAB (ALP+), FAP (PDGFRA+) and IC (ALP- PDFRA-) fractions from two separate microbiopsies obtained in parallel from the same subject (TD n = 2). **(B)** Full data-set on FACS fractions of CD56- ALP- PDGFRa+ population (BoNT naive n = 6, BoNT treated n = 9, TD n = 13). **(C)** Full data-set on FACS fractions of CD56- ALP- PDGFRa- population. (BoNT naive n = 6, BoNT treated n = 9, TD n = 13). **(D)** Linear regression of age and FACS fractions of CD56- ALP- PDGFRa+ sorted cells. Trend line and 95% confidence interval are shown. $R^2$ is 0.04378. Each dot represents a patient with CP (n = 11, open: BoNT naive; closed: with BoNT treatment history). **(E)** Linear regression of age and FACS fractions of CD56- ALP- PDGFRa- sorted cells. Trend line and 95% confidence interval are shown. $R^2$ is 0.1359. Each dot represents a patient with CP (n = 11, open: BoNT naive; closed: with BoNT treatment history).
